# A large and rich EEG dataset for modeling human visual object recognition

**DOI:** 10.1101/2022.03.15.484473

**Authors:** Alessandro T. Gifford, Kshitij Dwivedi, Gemma Roig, Radoslaw M. Cichy

## Abstract

The human brain achieves visual object recognition through multiple stages of nonlinear transformations operating at a millisecond scale. To predict and explain these rapid transformations, computational neuroscientists employ machine learning modeling techniques. However, state-of-the-art models require massive amounts of data to properly train, and to the present day there is a lack of vast brain datasets which extensively sample the temporal dynamics of visual object recognition. Here we collected a large and rich dataset of high temporal resolution EEG responses to images of objects on a natural background. This dataset includes 10 participants, each with 82,160 trials spanning 16,740 image conditions. Through computational modeling we established the quality of this dataset in five ways. First, we trained linearizing encoding models that successfully synthesized the EEG responses to arbitrary images. Second, we correctly identified the recorded EEG data image conditions in a zero-shot fashion, using EEG synthesized responses to hundreds of thousands of candidate image conditions. Third, we show that both the high number of conditions as well as the trial repetitions of the EEG dataset contribute to the trained models’ prediction accuracy. Fourth, we built encoding models whose predictions well generalize to novel participants. Fifth, we demonstrate full end-to-end training of randomly initialized DNNs that output M/EEG responses for arbitrary input images. We release this dataset as a tool to foster research in visual neuroscience and computer vision.

## Introduction

Visual object recognition is a complex cognitive function that is computationally solved in multiple nonlinear stages by the human brain (Marr, 1980; Goodale & Milner, 1992; Van Essen et al., 1992; Riesenhuber & Poggio, 1999; Ullman, 2000; Grill-Spector et al., 2001; Malach et al., 2002; Carandini et al., 2005). Through these stages information is transformed from representations of simple visual features such as oriented edges to representations of object categories (Tanaka, 1996; Logothetis & Sheinberg, 1996). To understand the principles of these transformations, computational neuroscientists build and employ mathematical models that predict the brain responses to arbitrary visual stimuli and explain their underlying neural mechanisms (Wu et al., 2006; Guest & Martin, 2021). The performance of these models benefits from training with large datasets: as an example, deep neural networks (DNNs) (Fukushima et al., 1982), the current state-of-the-art computational models of the visual brain (Yamins & DiCarlo, 2016; Cichy & Kaiser, 2019; Kietzmann et al., 2019a; Richards et al., 2019; Saxe et al., 2021), are trained on hundreds of thousands of different data points (Russakovsky et al., 2015). Yet, due to the difficulty of brain data acquisition, neuroscientific datasets usually comprise no more than a few thousand trials per participant and a limited number of conditions (Kay et al., 2008; Cichy et al., 2014; Horikawa & Kamitani, 2017).

To address the data hunger of current modeling goals, recently pioneering efforts have been taken to record large datasets of functional magnetic resonance imaging (fMRI) responses to images (Chang et al., 2019; Allen et al., 2021). However, while providing excellent spatial resolution, fMRI data lacks the temporal resolution to resolve neural dynamics at the level at which they occur. Since neurons communicate at millisecond scales, high temporal resolution neural data is a crucial component for building models of the visual brain (Thorpe et al., 1996; van de Nieuwenhuijzen et al., 2013; Cichy et al., 2014; Harel et al., 2016; Seeliger et al., 2017; Bankson et al., 2018; Dijkstra et al., 2018). Thus, in the present study we collected a large millisecond resolution electroencephalography (EEG) dataset of human brain responses to images of objects on a natural background. We extensively sampled 10 participants, each being presented with 16,740 image conditions repeated over 82,160 trials from the THINGS database (Hebart et al., 2019) by using a time-efficient rapid serial visual presentation (RSVP) paradigm (Intraub, 1981; Keysers et al., 2001; Grootswagers et al., 2019).

We then leveraged the unprecedented size and richness of our dataset to train and evaluate DNN-based linearizing and end-to-end encoding models (Wu et al., 2006; Kay et al., 2008; Naselaris et al., 2011; van Gerven, 2017; Seeliger et al., 2017; Kriegeskorte & Douglas, 2019; Seeliger et al., 2021; Khosla et al., 2021; Allen et al., 2021) that synthesize EEG responses to arbitrary images. The results showcase the quality of the dataset and its potential for computational modeling in five ways. First, the synthesized EEG data is strongly resemblant to its biological counterpart, with robust predictions even at single participants’ level. Second, we built zero-shot identification algorithms (Kay et al., 2008; Seeliger et al., 2017; Horikawa & Kamitani, 2017) that achieved high performance accuracies even when identifying among very large candidate image conditions set sizes: 81.3% for a set size of 200 candidate image conditions, 21.15% for a set size of 150,000 candidate image conditions, and extrapolated accuracy > 10% for a set size of 3,650,000 candidate image conditions, where chance ≤ 0.5%. Third, we show that both the high number of conditions as well as the trial repetitions of the dataset contribute to the trained models’ prediction accuracy. Fourth, we demonstrate that the encoding models’ predictions generalize to novel participants. Fifth, for the first time to our knowledge we demonstrate full end-to-end training (Seeliger et al., 2021; Khosla et al., 2021; Allen et al., 2021) of randomly initialized DNNs that output M/EEG responses for arbitrary input images.

We release the dataset as a tool to foster research in computational neuroscience and to bridge the gap between biological and artificial vision. We believe this will be of great use to further understanding of visual object recognition through the development of high-temporal resolution computational models of the visual brain, and to optimize artificial intelligence models through biological intelligence data (Sinz et al., 2019; Hassabis et al., 2017; Ullman, 2019; Toneva & Wehbe, 2019; Yang et al., 2022). Also all code used to generate the presented results accompanies the data release.

## Results

### A large and rich EEG dataset of visual responses to objects on a natural background

We used a RSVP paradigm (Intraub, 1981; Keysers et al., 2001; Grootswagers et al., 2019) to collect a large EEG dataset of visual responses to images of objects on a natural background (**Figure 1A**). This dataset contains data for 10 participants who viewed 16,540 training image conditions (**Figure 1B**) and 200 test image conditions (**Figure 1C**) coming from the THINGS database (Hebart et al., 2019). To allow for unbiased modeling the training and test images did not have any overlapping object concepts. We presented each training image condition 4 times and each test image condition 80 times, for a total of 82,160 image trials per participant over the course of four sessions. Thanks to the time-efficiency of the RSVP paradigm we collected up to 15 times more data than other typical recent M/EEG datasets used for modeling (Cichy et al., 2014; Seeliger et al., 2017). This allowed us to extensively sample single participants while drastically reducing the experimental time. During preprocessing we epoched the EEG recordings from −200ms to 800ms with respect to image onset, downsampled the resulting image epoch trials to 100 time points, and retained only the 17 occipital and parietal channels. As the basis of all further data assessment we aggregated the EEG recordings into a *biological training* (BioTrain) data matrix of shape (16,540 training image conditions × 4 condition repetitions × 17 EEG channels × 100 EEG time points) and a *biological test* (BioTest) data matrix of shape (200 test image conditions × 80 condition repetitions × 17 EEG channels × 100 EEG time points), for each participant. Providing this EEG data in its raw as well as preprocessed form is the major contribution of this resource.

**Figure 1.**
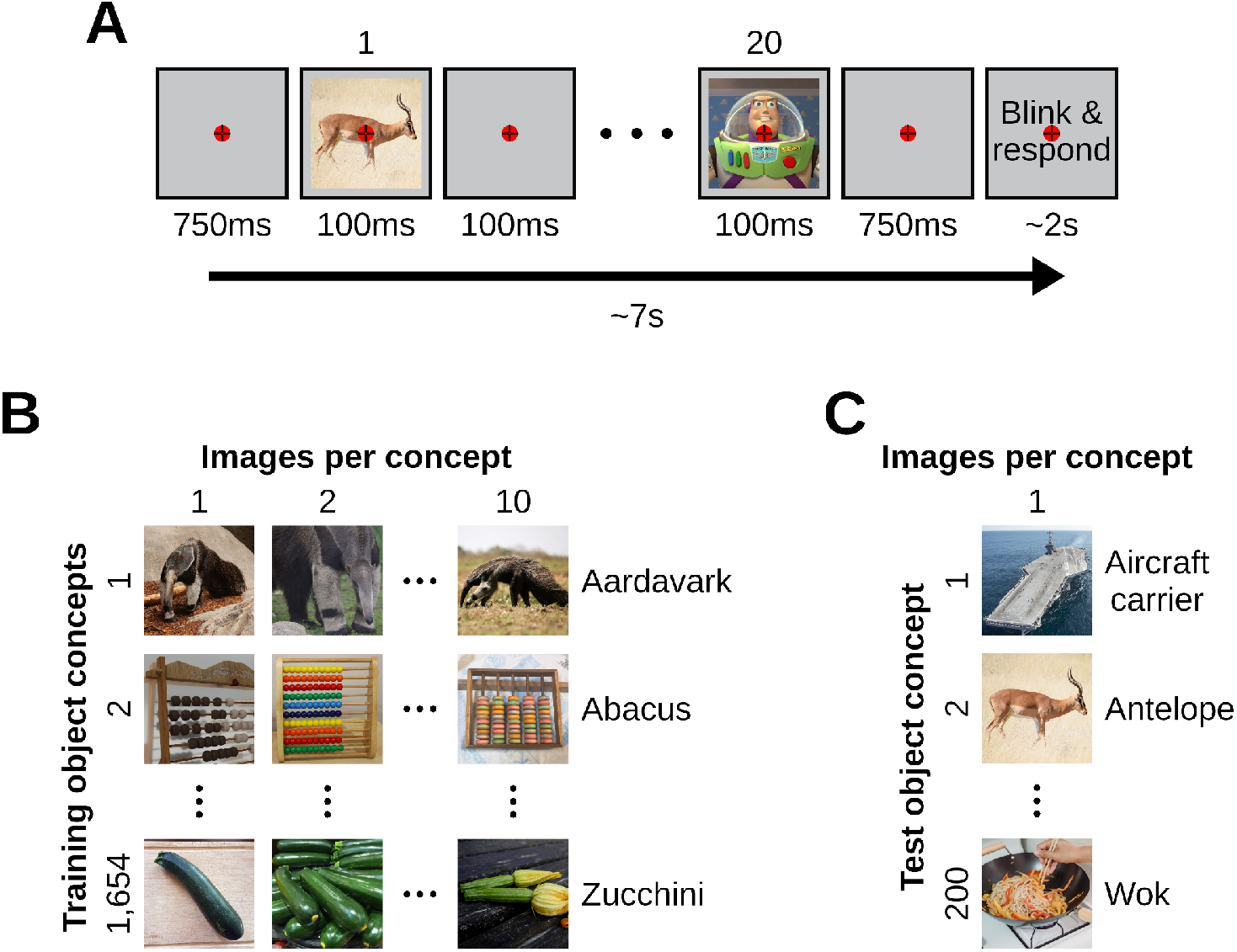
Experimental paradigm and stimuli images. (**A**) We presented participants with images of objects on a natural background using a RSVP paradigm. The paradigm consisted of rapid serial sequences of 20 images. Every sequence started with 750ms of blank screen, then each image was presented centrally for 100ms and a stimulus onset asynchrony (SOA) of 200ms, and it ended with another 750ms of blank screen. After every rapid sequence there were up to 2s during which we instructed participants to first blink and then report, with a keypress, whether the target image appeared in the sequence. We asked participants to gaze at a central bull’s eye fixation target present throughout the entire experiment. (**B**) The training image partition contains 1,654 object concepts of 10 images each, for a total of 16,540 image conditions. (**C**) The test image partition contains 200 object concepts of 1 image each, for a total of 200 image conditions.

### Building linearizing encoding models of EEG visual responses

We then assessed the suitability of this dataset for the development of computational models of the visual brain. We employed the training and test data, respectively, to build and evaluate linearizing encoding models which predict individual participant’s EEG visual responses to arbitrary images (Wu et al., 2006; Kay et al., 2008; Naselaris et al., 2011; van Gerven, 2017; Kriegeskorte & Douglas, 2019). We based our encoding algorithm on deep neural networks (DNNs), connectionist models which in the last decade have excelled in predicting human and non-human primate visual brain responses (Cadieu et al., 2014; Yamins et al., 2014; Güçlü & van Gerven, 2015; Storrs et al., 2021). The building of encoding models involved two steps. In the first step we non-linearly transformed the image pixel values using four DNNs pre-trained on ILSVRC-2012 (Russakovsky et al., 2015) commonly used for modeling brain responses: AlexNet (Krizhevsky, 2014), ResNet-50 (He et al, 2016), CORnet-S (Kubilius et al., 2019) and MoCo (Chen et al., 2020). Separately for each DNN we fed the training and test images, extracted the corresponding feature maps across all layers, appended the layers’ data together and downsampled it to 1,000 principal components using principal component analysis (PCA), resulting in the training DNN feature maps matrix of shape (16,540 training image conditions × 1,000 features) and the test DNN feature maps matrix of shape (200 test image conditions × 1000 features). In the second step we fitted the weights ***W***_***t,c***_ of several linear regressions that independently predicted each EEG feature’s response (i.e., the EEG activity at each combination of time points (***t***) and channels (***c***)) to the training images by linearly combining the training feature maps of each DNN (**Figure 2A**). We then multiplied the learned ***W***_***t,c***_ with the test DNN feature maps, obtaining the *synthetic test* (SynTest) EEG data matrix of shape (200 test image conditions × 17 EEG channels × 100 EEG time points) (**Figure 2B**). Following this procedure we obtained different instances of SynTest data for each participant and DNN.

**Figure 2.**
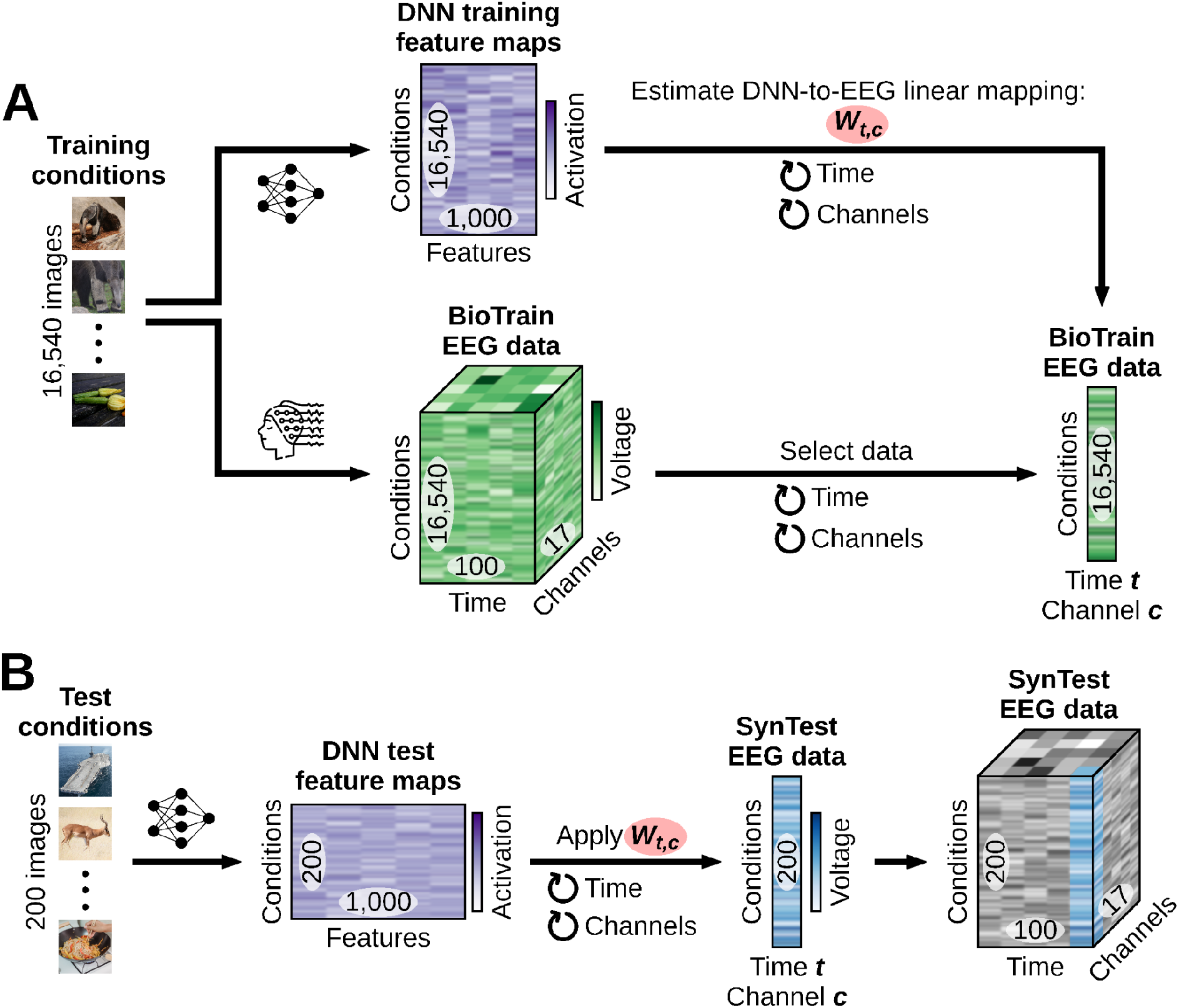
Linearizing encoding algorithm. For ease of visualization, here and in the following figures we omit the EEG condition repetitions dimension. (**A**) Through the training image conditions we obtained the training DNN feature maps and the BioTrain EEG data, and used them to build linearizing encoding models of EEG visual responses. For each combination of EEG features (time points (***t***) and channels (***c***)) we estimated the weights ***Wt,c*** of a linear regression using the corresponding single-feature BioTrain data as criterion and the training images DNN feature maps as predictors. (**B**) To obtain the SynTest EEG data we extracted the DNN feature maps of the test images, and multiplied them with the estimated ***Wt,c***.

### The BioTest EEG data is well predicted by linearizing encoding models

To evaluate the linearizing encoding models’ predictive power we quantified the similarity between the SynTest data and the BioTest data through a Pearson’s correlation (**Figure 3A**). We correlated each SynTest data EEG feature (i.e., each combination of EEG time points (***t***) and channels (***c***)) with the corresponding BioTest data feature (across the 200 test image conditions), resulting in a correlation coefficient matrix of shape (17 EEG channels × 100 EEG time points). We then averaged this matrix across the channels dimension, obtaining a correlation coefficient result vector with 100 components, one for each EEG time point.

**Figure 3.**
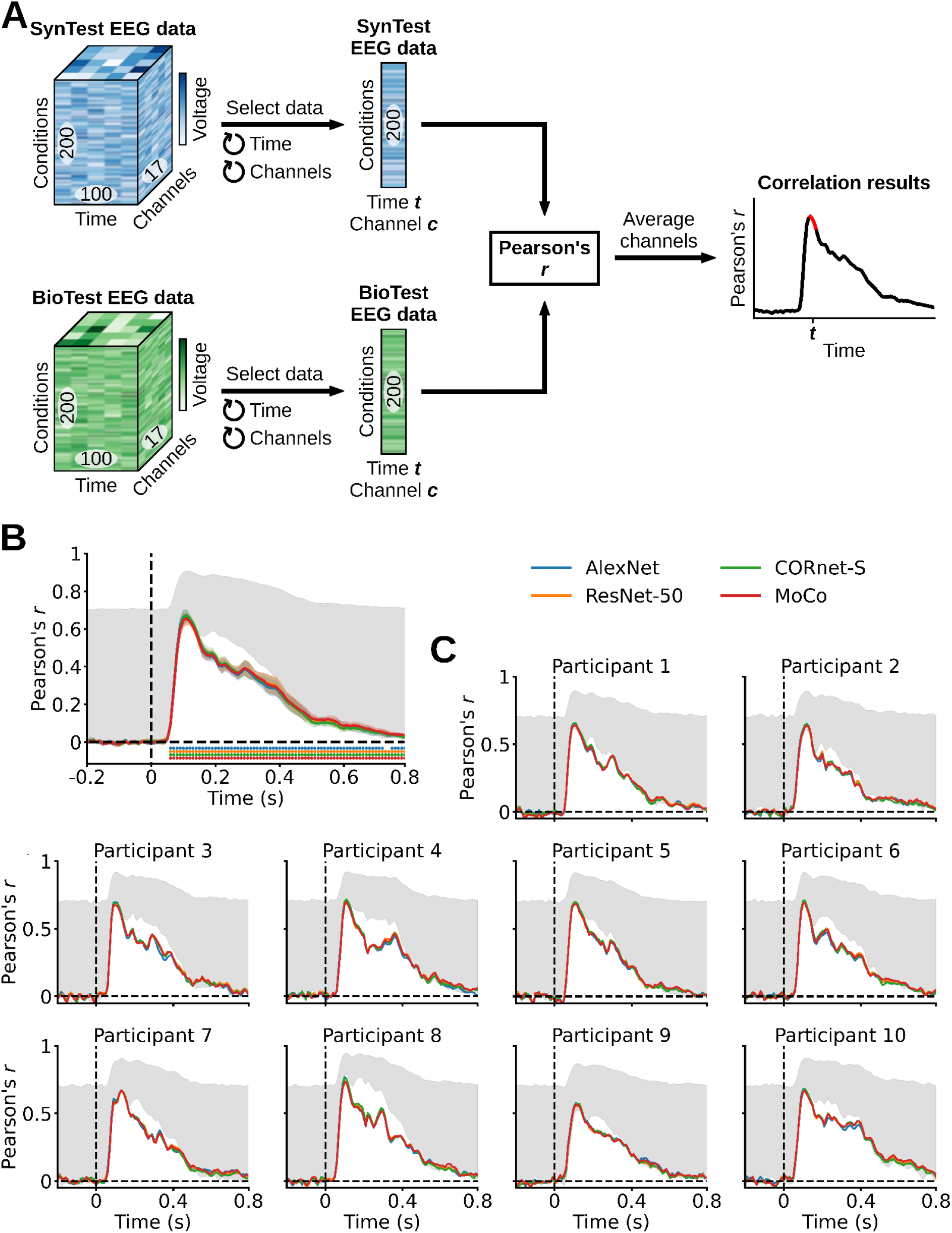
Evaluating the linearizing encoding models’ prediction accuracy through a correlation analysis. (**A**) We correlated each combination of SynTest EEG data features (time points (***t***) and channels (***c***)) with the corresponding combination of BioTest EEG data features, across the 200 test image conditions, and then averaged the correlation coefficients across channels. (**B**) Correlation results averaged across participants. The SynTest data is significantly correlated to the BioTest data from 60ms after stimulus onset until the end of the EEG epoch (*P* < 0.05, one-sample one-sided t-test, Bonferroni-corrected), with a peak at 110ms. (**C**) Individual participants’ results. Error margins reflect 95% confidence intervals. Rows of asterisks indicate significant time points (*P* < 0.05, one-sample one-sided t-tests, Bonferroni-corrected). In gray is the area between the noise ceiling lower and upper bounds, the black dashed vertical lines indicate onset of image presentation, and the black dashed horizontal lines indicate the chance level of no experimental effect.

As a complementary way to evaluate the linearizing encoding models’ predictive power we quantified the similarity between the SynTest data and the BioTest data through decoding (**Figure 4A**). Decoding is a commonly used method in computational neuroscience which exploits similar information present between the trials of each experimental condition to classify neural data (Haynes & Rees, 2006; Mur et al., 2009). If the SynTest data and the BioTest data have similar information, a decoding algorithm trained on the BioTest data would generalize its performance also to the SynTest data. We tested this through pairwise decoding: we trained linear support vector machines (SVMs) to perform binary classification between each pair of the 200 BioTest data image conditions, and then tested them on the corresponding pairs of SynTest data image conditions. We performed this analysis independently for each time point (***t***), resulting in a decoding accuracy matrix of shape (19,900 image condition pairs × 100 EEG time points). We then averaged this matrix across the image condition pairs dimension, obtaining a decoding accuracy result vector with 100 components, one for each EEG time point.

**Figure 4.**
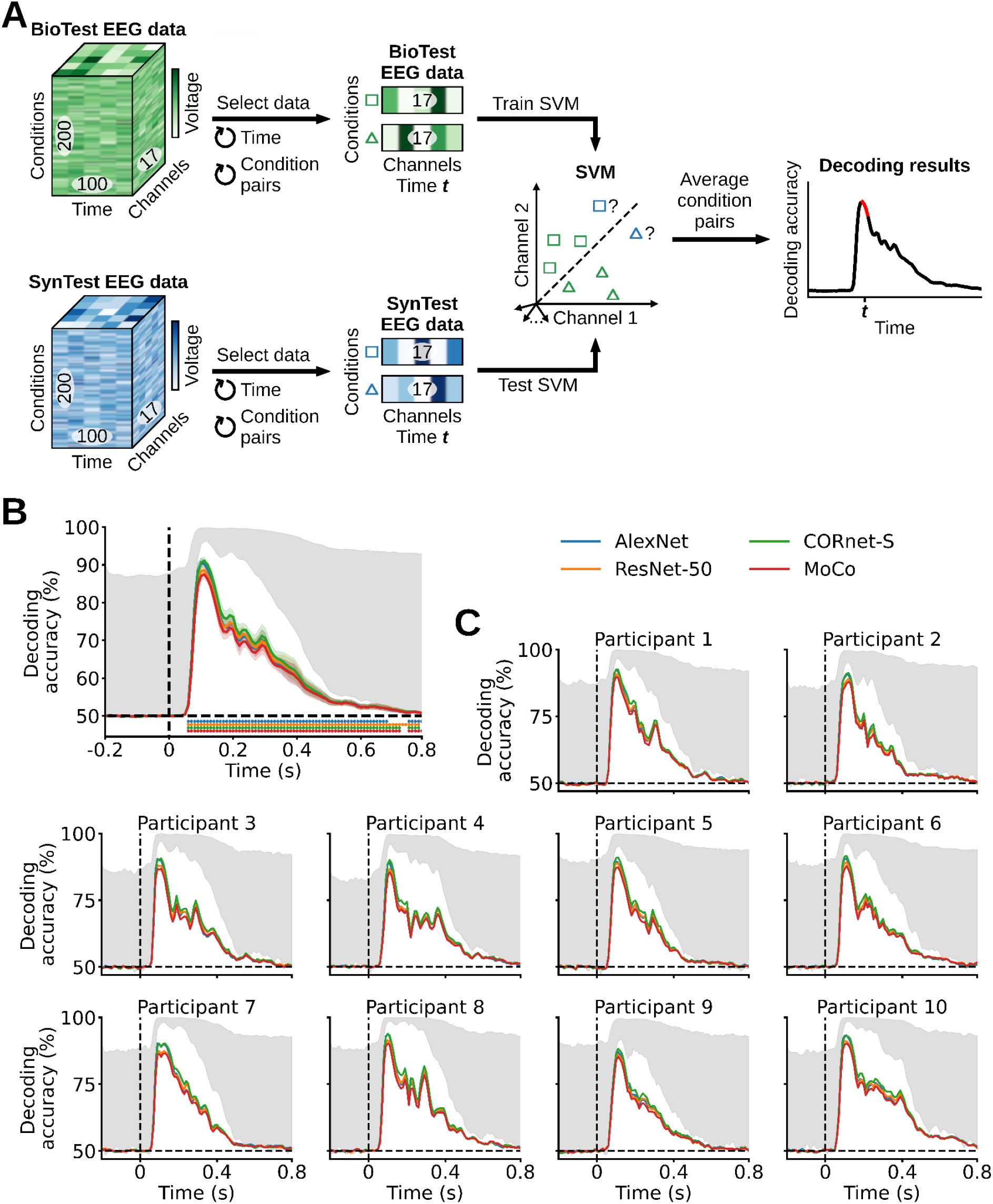
Evaluating the linearizing encoding models’ prediction accuracy through a pairwise decoding analysis. (**A**) At each time point (***t***) we trained an SVM to classify between two BioTest EEG data image conditions (using the channels vectors) and tested it on the two corresponding SynTest EEG data image conditions. We repeated this procedure across all image condition pairs, and then averaged the decoding accuracies across pairs. (**B**) Pairwise decoding results averaged across participants. The linear classifiers trained on the BioTest data significantly decode the SynTest data from 60ms after stimulus onset until the end of the EEG epoch (*P* < 0.05, one-sample one-sided t-test, Bonferroni-corrected), with peaks at 100-110ms. (**C**) Individual participants’ results. Error margins, asterisks, gray area and black dashed lines as in **Figure 3**.

We observe that the correlation results averaged across participants start being significant at 60ms after stimulus onset, and remain significantly above chance until the end of the EEG epoch at 800ms (*P* < 0.05, one-sample one-sided t-test, Bonferroni-corrected). Significant correlation peaks occur for all DNNs at 110ms after stimulus onset, with AlexNet, ResNet-50, CORnet-S and MoCo having correlation coefficients of, respectively, 0.67, 0.66, 0.67 and 0.66 (*P* < 0.05, one-sample one-sided t-test, Bonferroni-corrected), where the chance level is 0 (**Figure 3B**). Similarly, the pairwise decoding results averaged across participants start being significant at 60ms after stimulus onset, with significant effects present until the end of the EEG epoch at 800ms (*P* < 0.05, one-sample one-sided t-test, Bonferroni-corrected). Significant decoding peaks occur for all DNNs at 100-110ms after stimulus onset, with AlexNet, ResNet-50, CORnet-S and MoCo having decoding accuracies of, respectively, 90.37%, 88.57%, 91.06% and 87.45% (*P* < 0.05, one-sample one-sided t-test, Bonferroni-corrected), where the chance level is 50% (**Figure 4B**). All participants yielded qualitatively similar results (**Figure 3C**, **Figure 4C**). Taken together, these results show that the linearizing encoding models are successful in predicting EEG data which robustly and significantly resembles its biological counterpart. Further, they show that each participant’s neural responses can be consistently predicted in isolation, thus highlighting the quality of the visual information contained in our EEG dataset and its potential for the development of new high-temporal resolution models and theories of the visual brain.

### The BioTest data is significantly identified in a zero-shot fashion using synthesized data of up to 150,200 candidate images

The previous analyses showed that our linearizing encoding models synthesize EEG data that significantly resembles its biological counterpart. Here we explored whether we can leverage this high prediction accuracy to build algorithms that identify the image conditions of the BioTest data in a zero-shot fashion, namely, that identify arbitrary image conditions without prior training. If possible, this would contribute to the goal of building models capable of identifying potentially infinite neural data conditions on which they were never trained (Kay et al., 2008; Seeliger et al., 2017; Horikawa & Kamitani, 2017) (**Figure 5A**). For the identification we used the SynTest and the *synthetic Imagenet* (SynImagenet) data, where the latter consisted of the synthesized EEG responses to the 150,000 validation and test images coming from the ILSVRC-2012 image set (Russakovsky et al., 2015), organized in a data matrix of shape (150,000 image conditions × 17 EEG channels × 100 EEG time points). Importantly, those images did not overlap with either the image set for which EEG data was recorded. The further analysis involved two steps: feature selection and identification.

**Figure 5.**
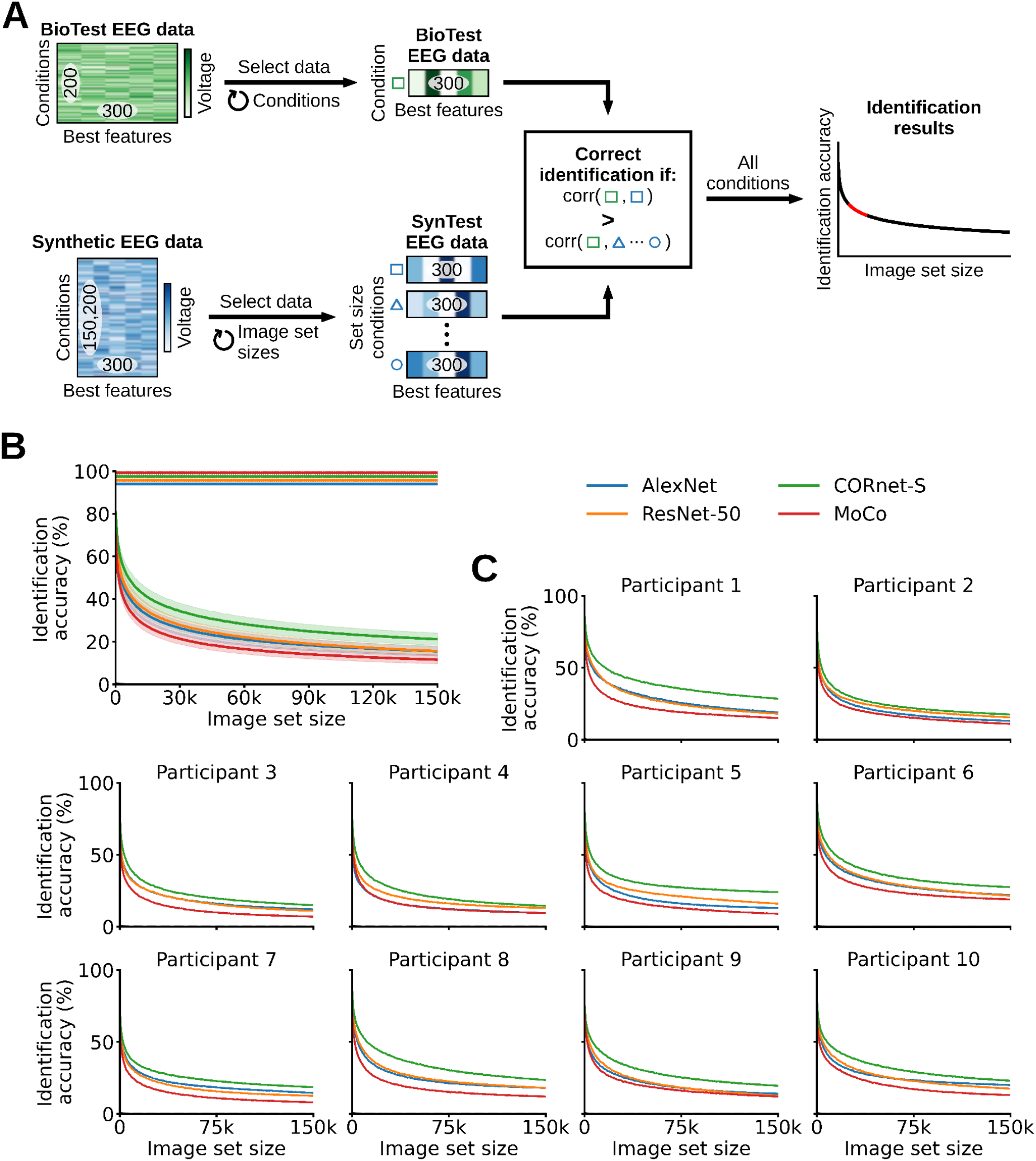
Zero-shot identification of the BioTest data using the SynTest data and the synthesized EEG visual responses to the 150,000 ILSVRC-2012 validation and test image conditions (SynImagenet). (**A**) We correlated the best features of each BioTest data condition with different image set sizes of candidate synthetic image conditions (SynTest + SynImagenet data). At each image set size, a BioTest data condition is correctly identified if it is mostly correlated to its corresponding SynTest data condition, among all other synthetic data conditions. (**B**) Zero-shot identification results averaged across participants. With a SynImagenet set size of 0 the synthesized data of AlexNet, ResNet-50, CORnet-S, MoCo significantly identify the BioTest data with accuracies of, respectively, 75.05%, 75.85%, 81.3%, 70.9%. (*P* < 0.05, one-sample one-sided t-test, Bonferroni-corrected). With a SynImagenet set size of 150,000 the synthesized data of AlexNet, ResNet-50, CORnet-S, MoCo significantly identify the BioTest data with accuracies of, respectively, 15.5%, 15.55%, 21.15%, 11.55%. (**C**) Individual participants’ results. Rows of asterisks indicate significant image set sizes (*P* < 0.05, one-sample one-sided t-tests, Bonferroni-corrected). Error margins and black dashed lines as in **Figure 3**.

In the feature selection step we retained the 300 EEG channels and time points best predicted by the encoding models, as narrowing down the EEG data to these features improved the identification accuracy. In detail, we synthesized the EEG responses to the 16,540 training images, obtaining the *synthetic train* (SynTrain) data matrix of shape (16,540 training image conditions × 17 EEG channels × 100 EEG time points). We then correlated each BioTrain data feature (i.e., each combination of EEG channels and EEG time points) with the corresponding SynTrain data feature (across the 16,540 training image conditions), and only retained the 300 SynTest, BioTest and SynImagenet data EEG features corresponding to the 300 highest correlation scores. This resulted in feature vectors of 300 components for each image condition.

In the identification step we correlated the feature vectors of each BioTest data image condition with the feature vectors of all the candidate image conditions, where the candidate image conditions corresponded to the SynTest data image conditions plus a varying amount of SynImagenet data image conditions. We increased the set sizes of the SynImagenet candidate image conditions from 0 to 150,000 with steps of 1,000 images (for a total of 151 set sizes), and performed the identification at every set size. At each set size a BioTest data image condition is considered correctly identified if the correlation coefficient between its feature vector and the feature vector of the corresponding SynTest data image condition is higher than the correlation coefficients between its feature vector and the feature vectors of all other candidate image conditions. We calculated identification accuracies through the ratio of successfully decoded image conditions over all 200 BioTest image conditions, obtaining a zero-shot identification result vector with 151 components, one for each candidate image set size. The results of the correct SynTest data image condition falling within the three or ten most correlated image conditions can be seen in **Supplementary Figures 3-4** and **Supplementary Tables 1-2**.

The zero-shot identification results averaged across participants are significant for all SynImagenet set sizes (*P* < 0.05, one-sample one-sided t-test, Bonferroni-corrected). With a SynImagenet set size of 0 (corresponding to using only the 200 SynTest data image conditions as candidate image conditions) the BioTest data image conditions are identified by AlexNet, ResNet-50, CORnet-S and MoCo with accuracies of, respectively, 75.05%, 75.85%, 81.3%, 70.9%, where the chance level is equal to 1 / 200 test image conditions = 0.5%. As the SynImagenet set size increases the identification accuracies monotonically decrease. With a SynImagenet set size of 150,000 (corresponding to using the 200 SynTest data plus the 150,000 SynImagenet data image conditions as candidate image conditions) the BioTest data image conditions are identified by AlexNet, ResNet-50, CORnet-S and MoCo with accuracies of, respectively, 15.5%, 15.55%, 21.15%, 11.55%, where the chance level is equal to 1 / (200 test image conditions + 150,000 ILSVRC-2012 image conditions) < 10^−5^% (**Figure 5B**). To extrapolate the identification accuracies to potentially larger candidate image set sizes we fit a power-law function to the results. We averaged the extrapolations across participants, and found that the identification accuracy would remain above 10% with a candidate image set size of 914,000 for AlexNet, 588,000 for ResNet-50, 3,650,000 for CORnet-S and 348,000 for MoCo, and above 0.5% (the original chance level) with a candidate image set size of 2.18E+11 for AlexNet, 3.43E+09 for ResNet-50, 1.62E+13 for CORnet-S and 1.11E+10 for MoCo (**Table 1**). All participants yielded qualitatively similar results (**Figures 5C; Table 1**). These results demonstrate that our dataset allows building algorithms that reliably identify arbitrary neural data conditions, in a zero-shot fashion, among millions of possible alternatives.

**Table 1.** Extrapolation of the zero-shot identification accuracy as a function of candidate image set sizes. The values in the table indicate the candidate image set sizes required for the identification accuracy to drop below 10% and 0.5%.

|  | Identification accuracy < 10% |  |  |  | Identification accuracy < 0.5% |  |  |  |
| --- | --- | --- | --- | --- | --- | --- | --- | --- |
|  | AlexNet | ResNet-50 | CORnet-S | MoCo | AlexNet | ResNet-50 | CORnet-S | MoCo |
| Participant 1 | 7.35E+05 | 6.31E+05 | 5.68E+06 | 5.30E+05 | 1.10E+09 | 8.44E+08 | 1.66E+11 | 4.56E+09 |
| Participant 2 | 2.97E+05 | 5.38E+05 | 9.02E+05 | 1.93E+05 | 7.27E+08 | 2.76E+09 | 1.24E+10 | 1.96E+08 |
| Participant 3 | 2.35E+05 | 1.79E+05 | 4.38E+05 | 7.31E+04 | 3.53E+08 | 7.34E+07 | 1.12E+09 | 2.56E+07 |
| Participant 4 | 1.33E+05 | 3.28E+05 | 3.67E+05 | 1.27E+05 | 4.70E+08 | 3.22E+09 | 5.38E+08 | 5.18E+08 |
| Participant 5 | 3.12E+05 | 6.15E+05 | 1.65E+07 | 1.24E+05 | 2.27E+09 | 3.22E+09 | 1.61E+14 | 6.03E+07 |
| Participant 6 | 2.01E+06 | 1.41E+06 | 7.06E+06 | 1.58E+06 | 3.91E+10 | 7.95E+09 | 6.80E+11 | 1.03E+11 |
| Participant 7 | 5.27E+05 | 2.80E+05 | 1.25E+06 | 9.11E+04 | 8.31E+09 | 1.80E+09 | 3.54E+10 | 6.71E+07 |
| Participant 8 | 1.27E+06 | 9.67E+05 | 1.63E+06 | 2.50E+05 | 8.13E+10 | 1.20E+10 | 6.64E+09 | 1.23E+09 |
| Participant 9 | 3.50E+05 | 2.46E+05 | 9.08E+05 | 2.39E+05 | 8.76E+08 | 9.52E+07 | 2.83E+09 | 3.40E+08 |
| Participant 10 | 3.28E+06 | 6.93E+05 | 1.84E+06 | 2.71E+05 | 2.04E+12 | 2.31E+09 | 1.46E+10 | 3.17E+08 |
| Average | 9.14E+05 | 5.88E+05 | 3.65E+06 | 3.48E+05 | 2.18E+11 | 3.43E+09 | 1.62E+13 | 1.11E+10 |

### The amount of training image conditions and condition repetitions both contribute to modeling quality

To understand which aspects of our EEG dataset contribute to its successful modeling we examined the linearizing encoding models’ prediction accuracy as a function of the amount of trials with which they are trained. The amount of training trials is determined by two factors: the number of image conditions and the number of EEG repetitions per each image condition. Both factors may improve the modeling of neural responses in different ways, as high numbers of image conditions lead to a richer training set which more comprehensively samples the representational space underlying vision, and high numbers of condition repetitions increase the signal to noise ratio (SNR) of the training set.

To disentangle the effect of both factors we trained linearizing encoding models using different quartiles of training image conditions (4,135, 8,270, 12,405, 16,540) and condition repetitions (1, 2, 3, 4), and tested their predictions through the correlation analysis. We performed an ANOVA on the correlation results averaged over participants, EEG features (all channels; time points between 60-500ms) and DNN models, and observed a significant effect of both number of image conditions and condition repetitions, along with a significant interaction of the two factors (*P* < 0.05, two-way repeated measures ANOVA) (**Figure 6A**). All participants yielded qualitatively similar results (**Supplementary Figure 5**). This suggests that the amount of image conditions and condition repetitions both improve the modeling of neural data.

**Figure 6.**
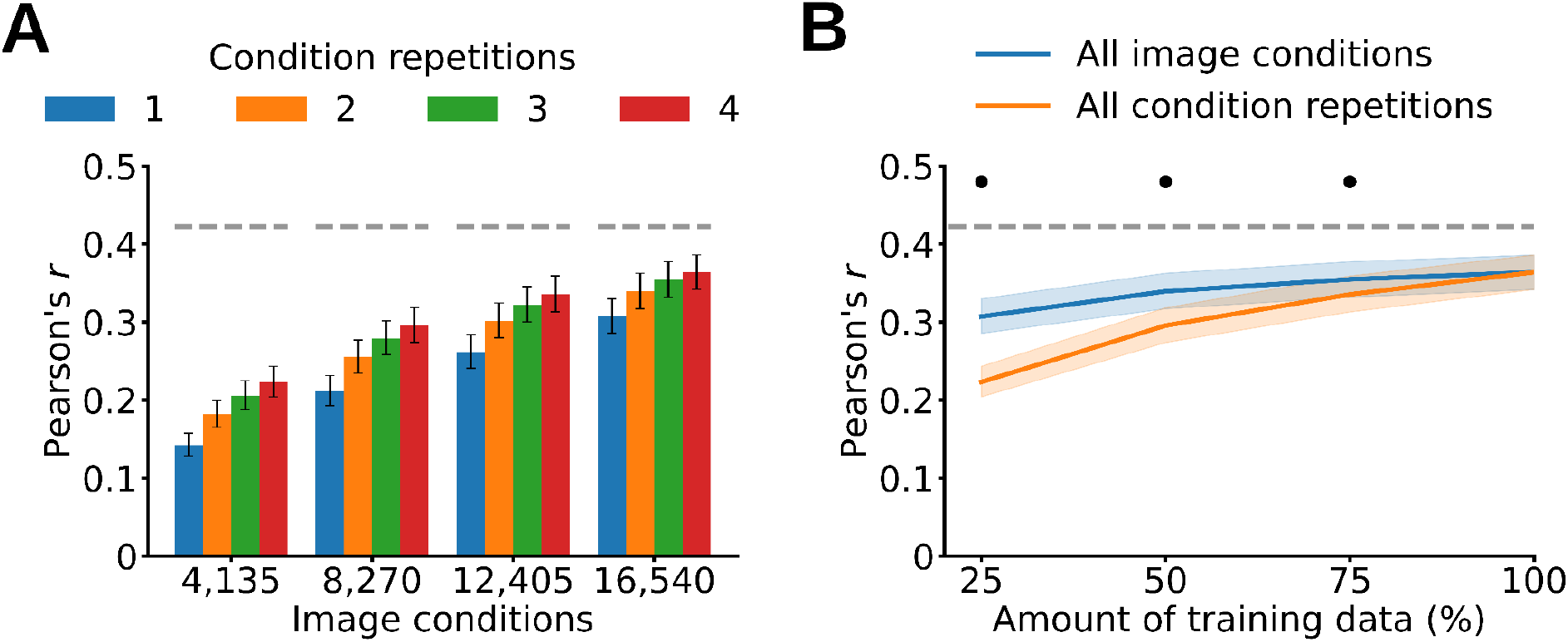
Linearizing encoding models’ prediction accuracy as a function of training data. (**A**) Training linearizing encoding models using different quartiles of image conditions and condition repetitions result in a significant effect of both factors (*P* < 0.05, two-way repeated measures ANOVA). (**B**) Training linearizing encoding models using all image conditions leads to higher prediction accuracies than training them using all condition repetitions (*P* < 0.05, repeated measures two-sided t-test, Bonferroni-corrected). The gray dashed line represents the noise ceiling lower bound. The asterisks indicate a significant difference between all image conditions and all condition repetitions (*P* < 0.05, repeated measures two-sided t-test, Bonferroni-corrected). Error margins and gray dashed lines as in **Figure 3**.

We then asked which of the two factors contributes more to the linearizing encoding models’ prediction accuracy. For this we compared model prediction accuracy for cases where the number of repetitions or conditions differed, but the total number of trials was the same. As we had four trial repetitions, we divided the total amount of training trials into quartiles (25%, 50%, 75% and 100% of the total training trials). At each quartile we trained linearizing encoding models using all image conditions and the quartile’s percentage of condition repetitions, and tested their predictions through the correlation analysis. For example, at the first quartile we trained linearizing encoding models using all image conditions and one condition repetition, corresponding to 25% of the total training data. To compare, we repeated the same procedure while using all condition repetitions and the quartile’s percentage of image conditions. The correlation results averaged across participants, EEG features (all channels; time points between 60-500ms) and DNNs show that using all image conditions (and quartiles of condition repetitions) leads to higher prediction accuracies than using all condition repetitions (and quartiles of image conditions) (*P* < 0.05, repeated measures two-sided t-test, Bonferroni-corrected) (**Figure 6B**). All participants yielded qualitatively similar results (**Supplementary Figure 6**). This indicates that although both factors improve the modeling of neural data, the amount of image conditions does so here to a larger extent.

### The linearizing encoding models’ predictions generalize across participants

Next we explored whether our linearizing encoding models’ predictions generalize to new participants. We asked: Can we accurately synthesize a participant’s EEG responses without using any of their data for the encoding models’ training? If possible, our dataset could serve as a useful benchmark for the development and assessment of methods that combine EEG data across participants (Haxby et al., 2020; Richard et al., 2020; Kwon et al., 2019; Zhang et al., 2021). To verify this we trained linearizing encoding models on the averaged SynTrain EEG data of all minus one participants (**Figure 7A**), and tested their predictions against the BioTest data of the left out participant through the correlation and pairwise decoding analyses (**Figure 7B**). We repeated this procedure for all participants.

**Figure 7.**
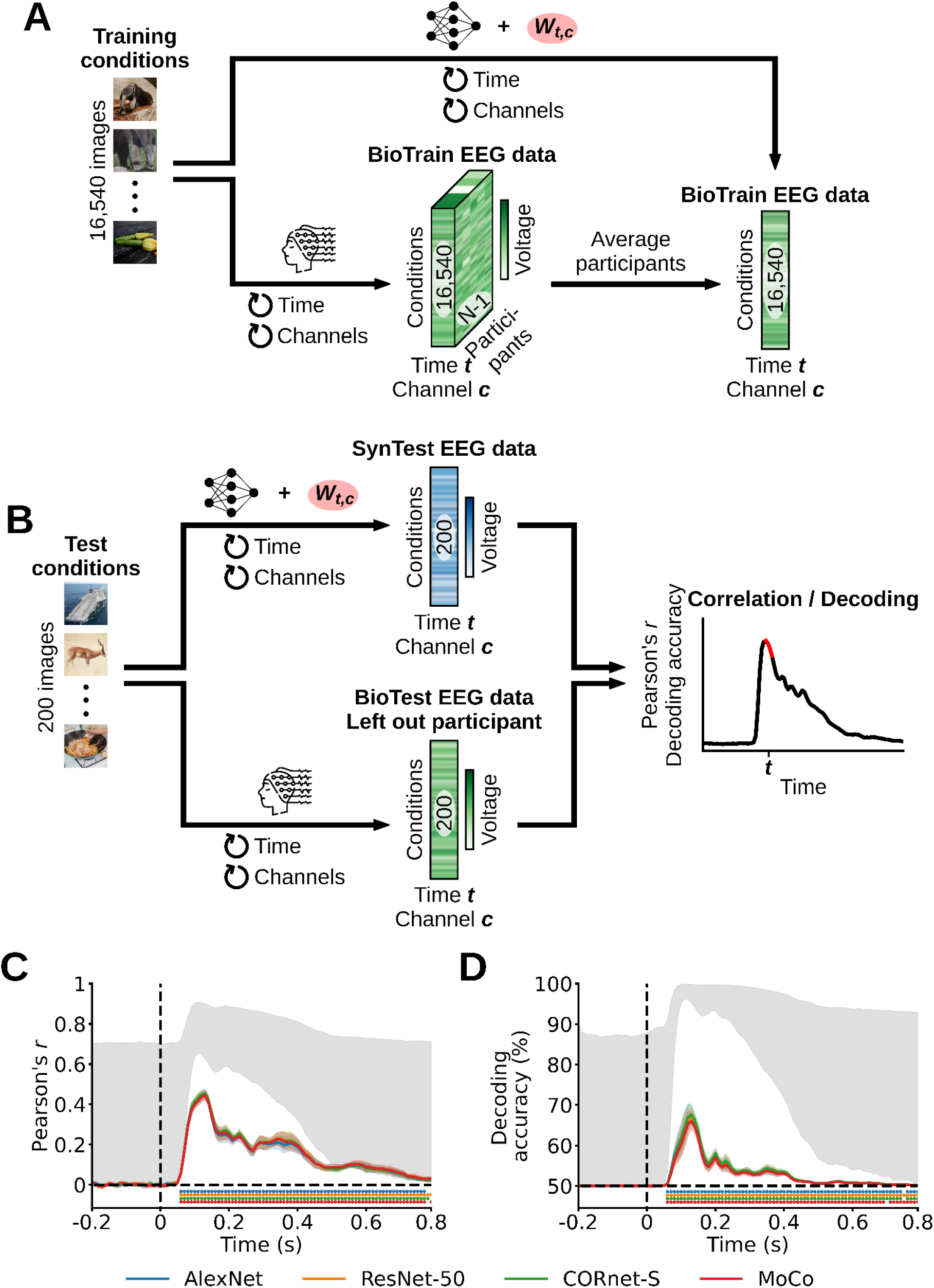
Evaluating the prediction accuracy of linearizing encoding models which generalize to novel participants, through correlation and pairwise decoding analyses. (**A**) We trained linearizing encoding models on the averaged SynTrain EEG data of all minus one participants. (**B**) We tested the encoding models’ predictions against the BioTest data of the left out participant through the correlation and pairwise decoding analyses. (**C**) Correlation results averaged across participants. The SynTest data is significantly correlated to the BioTest data from 60ms after stimulus onset until the end of the EEG epoch (*P* < 0.05, one-sample one-sided t-test, Bonferroni-corrected), with a peak at 130ms. (**D**) Pairwise decoding results averaged across participants. The linear classifiers trained on the BioTest data significantly decode the SynTest data from 60ms after stimulus onset until the end of the EEG epoch (*P* < 0.05, one-sample one-sided t-test, Bonferroni-corrected), with a peak at 130ms. Error margins, asterisks, gray area and black dashed lines as in **Figure 3**.

When averaging the Pearson correlation coefficients across participants we observe that the correlation between the SynTest data and the BioTest data starts being significant at 60ms after stimulus onset, and remains significantly above chance until the end of the EEG epoch at 800ms (*P* < 0.05, one-sample one-sided t-test, Bonferroni-corrected). Significant correlation peaks occur for all DNNs at 130ms after stimulus onset, with AlexNet, ResNet-50, CORnet-S and MoCo having correlation coefficients of, respectively, 0.45, 0.46, 0.46, 0.44 (*P* < 0.05, one-sample one-sided t-test, Bonferroni-corrected), where the chance level is 0 (**Figure 7C**). Likewise, the decoding accuracies averaged across participants start being significant at 60ms after stimulus onset, with significant effects present until the end of the EEG epoch at 800ms (*P* < 0.05, one-sample one-sided t-test, Bonferroni-corrected). Significant decoding peaks occur for all DNNs at 130ms after stimulus onset, with AlexNet, ResNet-50, CORnet-S and MoCo having decoding accuracies of, respectively, 67.44%, 66.62%, 67.63%, 66.01% (*P* < 0.05, one-sample one-sided t-test, Bonferroni-corrected), where the chance level is 50% (**Figure 7D**). In both analyses all participants yielded qualitatively similar results (**Supplementary Figures 7-8**). This shows that our EEG dataset is a suitable testing ground for methods which generalize and combine EEG data across participants.

### The BioTest EEG data is successfully predicted by end-to-end encoding models based on the AlexNet architecture

So far we predicted the synthetic data through the linearizing encoding framework, which relied on DNNs pre-trained on an image classification task. An alternative encoding approach, named end-to-end encoding (Seeliger et al., 2021; Khosla et al., 2021; Allen et al., 2021), is based on DNNs trained from scratch to predict the neural responses to arbitrary images. This direct infusion of brain data during the model’s learning could lead to DNNs having internal representations that more closely match the properties of the visual brain (Sinz et al., 2019; Allen et al., 2021). However, with a few exceptions (Seeliger et al., 2021; Khosla et al., 2021; Allen et al., 2021), the development of end-to-end encoding models has been so far prohibitive due to the large amount of data needed to train a DNN in combination with the small size of existing brain datasets. Thus, in this final analysis we exploited the largeness and richness of our EEG dataset to train randomly initialized AlexNet architectures to synthesize the EEG responses to images, independently for each participant. We started by replacing AlexNet’s 1000-neurons output layer with a 17-neurons layer, where each neuron corresponded to one of the 17 EEG channels. Then, for each EEG time point (***t***) we trained one such model to predict the multi-channel EEG responses to visual stimuli using the 16,540 training images as inputs and the corresponding BioTrain data as output targets (**Figure 8A**). Finally, we deployed the trained networks to synthesize the EEG responses to the 200 test images and evaluated their prediction accuracy through the same correlation and pairwise decoding analyses described above (**Figure 8B**).

**Figure 8.**
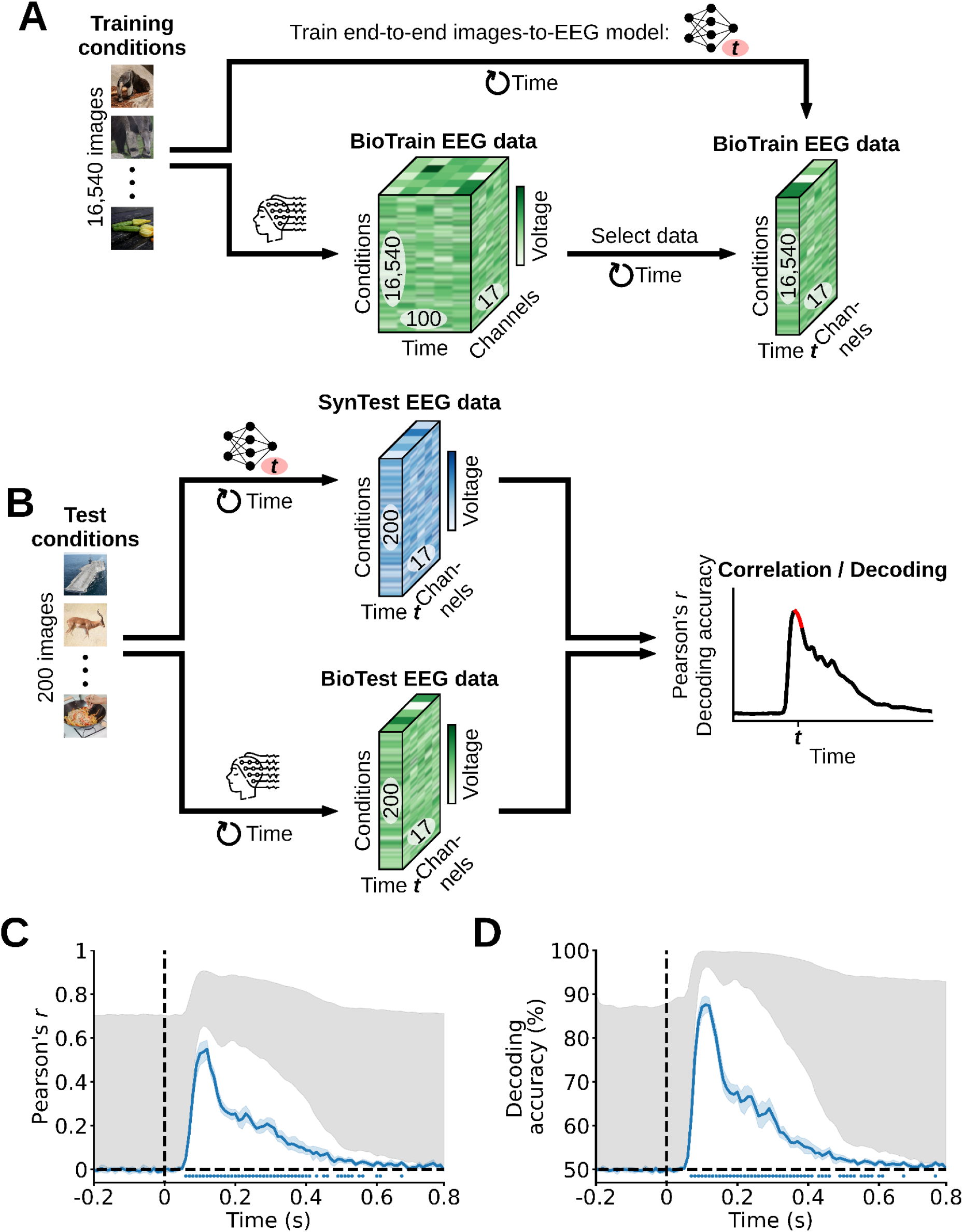
Evaluating the end-to-end encoding models’ prediction accuracy through correlation and pairwise decoding analyses. (**A**) At each EEG time point (***t***) we trained one encoding model end-to-end to predict the SynTrain data channel responses using the corresponding training images as input. (**B**) We used the trained encoding models to predict the SynTest data using the test images as input, and evaluated their prediction accuracies by comparing the SynTest and BioTest data through correlation and pairwise decoding analyses. (**C**) Correlation results averaged across participants. The SynTest data is significantly correlated to the BioTest data from 60ms after stimulus onset until 670ms (*P* < 0.05, one-sample one-sided t-test, Bonferroni-corrected), with a peak at 120ms. (**D**) Pairwise decoding results averaged across participants. The linear classifiers trained on the BioTest data significantly decode the SynTest data from 70ms after stimulus onset until 760ms (*P* < 0.05, one-sample one-sided t-test, Bonferroni-corrected), with peaks at 110ms. Error margins, asterisks, gray area and black dashed lines as in **Figure 3**.

We observe that the correlation results averaged across participants start being significant at 60ms after stimulus onset, with a correlation coefficient peak at 120ms of 0.55, and have significant effects until 670ms (*P* < 0.05, one-sample one-sided t-test, Bonferroni-corrected) (**Figure 8C**). Similarly, the pairwise decoding results averaged across participants start being significant at 70ms after stimulus onset, with a decoding accuracy peak at 110ms of 87.59%, and have significant effects until 760ms (*P* < 0.05, one-sample one-sided t-test, Bonferroni-corrected) (**Figure 8D**). All participants yielded qualitatively similar results (**Supplementary Figures 9-10**). This proves that our EEG dataset allows for the successful training of DNNs in an end-to-end fashion, paving the way for a stronger symbiosis between brain data and deep learning models benefitting both neuroscientists interested in building better models of the brain (Seeliger et al., 2021; Khosla et al., 2021; Allen et al., 2021) and computer scientists interested in creating better performing and more brain-like artificial intelligence algorithms through inductive biases from biological intelligence (Sinz et al., 2019; Hassabis et al., 2017; Ullman, 2019; Toneva & Wehbe, 2019; Yang et al., 2022).

## Discussion

### Summary

We used a RSVP paradigm (Intraub, 1981; Keysers et al., 2001; Grootswagers et al., 2019) to collect a large and rich EEG dataset of neural responses to images of real-world objects on a natural background, which we release as a tool to foster research in vision neuroscience and computer vision. Through computational modeling we established the quality of this dataset in five ways. First, we trained linearizing encoding models (Wu et al., 2006; Kay et al., 2008; Naselaris et al., 2011; van Gerven, 2017; Kriegeskorte & Douglas, 2019) that successfully synthesized the EEG responses to arbitrary images. Second, we correctly identified the recorded EEG data image conditions in a zero-shot fashion (Kay et al., 2008; Seeliger et al., 2017; Horikawa & Kamitani, 2017), using EEG synthesized responses to hundreds of thousands of candidate image conditions. Third, we show that both the high number of conditions as well as the trial repetitions of the EEG dataset contribute to the trained model’s prediction accuracy. Fourth, we built encoding models whose predictions well generalize to novel participants. Fifth, we demonstrate full end-to-end training (Seeliger et al., 2021; Khosla et al., 2021; Allen et al., 2021) of randomly initialized DNNs that output M/EEG responses for arbitrary input images.

### Size matters

In the last years cognitive neuroscientists have drastically increased the scope of their recordings from datasets with dozens of stimuli to datasets comprising several thousands of stimuli per participant (Chang et al., 2019; Naselaris et al., 2021; Allen et al., 2021). Compared to their predecessors, these large datasets more comprehensively sample the visual space and interact better with modern data-hungry machine learning algorithms. In this spirit we extensively sampled 10 participants with 82,160 trials spanning 16,740 image conditions, and showed how this unprecedented size contributes to high modeling performances. We released the data in both its raw and preprocessed format ready for modeling to allow researchers of different analytical perspectives to use the dataset in their preferred way immediately. We believe the largeness of this dataset holds great promise for neuroscientists interested in further improving theories and models of the visual brain, as well as computer scientists interested in improving machine vision models through biological vision constraints (Haxby et al., 2020; Richard et al., 2020; Kwon et al., 2019; Zhang et al., 2021).

### Linearizing encoding modeling

We showcased the potential of the dataset for modeling visual responses by building linearizing encoding algorithms (Wu et al., 2006; Kay et al., 2008; Naselaris et al., 2011; van Gerven, 2017; Kriegeskorte & Douglas, 2019) that predicted EEG visual responses to arbitrary images. Through correlation and decoding analyses we showed that the encoding models synthesized data which significantly resembles its biological counterpart robustly and consistently across all participants. These results highlight the signal quality of the visual information present in the EEG dataset, making it a promising candidate for the development of new high-temporal resolution models and theories of the neural dynamics of vision capable of predicting, decoding and even explaining visual object recognition.

### Zero-shot identification

Decoding models in neuroscience typically classify between only a few data conditions, while relying on data exemplars from these same conditions to train (Haynes & Rees, 2006; Mur et al., 2009). As a result, their performance fails to generalize to the unlimited space of different brain states. Here we exploited the prediction accuracy of the synthesized EEG responses to build zero-shot identification algorithms that identify potentially infinite neural data image conditions, without the need of prior training (Kay et al., 2008; Seeliger et al., 2017; Horikawa & Kamitani, 2017). Through this framework we identified the BioTest EEG image conditions between hundreds of thousands of candidate image conditions. Even when the identification algorithm failed to assign the correct image condition to the biological EEG responses, we show that it nevertheless selected a considerable amount (up to 45%) of the correct image conditions as the first three or ten choices (**Supplementary Figures 3-4**). These results suggest that our dataset is a good starting ground for the future creation of zero-shot identification algorithms to be deployed not only in research, but also in cutting-edge brain computer interface (BCI) technology (Abiri et al., 2019; Petit et al., 2021).

### Both number of image conditions and condition repetitions determine dataset quality

Building linearizing encoding algorithms with different amounts of training data revealed that the encoding models’ prediction accuracies are significantly affected by both the amount of EEG image conditions (to a higher extent) and repetitions of measurements (to a lower extent). Given that the noise ceiling lower bound estimate is not reached, these findings suggest that the prediction accuracy of the linearizing encoding models would have benefited from either more training data trials, or from a training dataset with the same amount of trials but having more image conditions and less repetitions of measurements. Based on these observations, for future dataset collections we recommend prioritizing the amount of stimuli conditions over the amount of repetitions of measurements.

### Inter-participant predictions

Typically, computational models in neuroscience are trained and evaluated on the data of single participants (Kay et al., 2008; Yamins et al., 2014; Güçlü & van Gerven, 2015; Seeliger et al., 2017; Horikawa & Kamitani, 2017). While this approach is well motivated by the neural idiosyncrasies of every individual (Charest et al., 2014), it fails to produce models that leverage the shared information across multiple brains. Here we show that our encoding models well predict out-of-set participants, indicating that our dataset is a suitable testing ground for methods which generalize and combine neural data across participants, as well as for BCI technology that can be readily used on novel participants without the need of calibration (Haxby et al., 2020; Richard et al., 2020; Kwon et al., 2019; Zhang et al., 2021).

### End-to-end encoding

So far limitations in neural dataset sizes led computational neuroscientists to model brain data mostly using pre-trained DNNs (Cadieu et al., 2014; Yamins et al., 2014; Güçlü & van Gerven, 2015; Naselaris et al., 2015; Seeliger et al., 2017). Here, we leveraged the largeness and richness of our dataset to demonstrate, for the first time to our knowledge with EEG data, the feasibility of training a randomly initialized AlexNet architecture to predict the neural responses to arbitrary images in an end-to-end fashion (Seeliger et al., 2021; Khosla et al., 2021; Allen et al., 2021). The end-to-end approach opens the doors to training complex computational algorithms directly with brain data, potentially leading to models which more closely mimic the internal representation of the visual system (Sinz et al., 2019; Allen et al., 2021). This will in turn make it possible for computer scientists to use the neural representations of biological systems as inductive biases to improve artificial systems under the assumption that increasing the brain-likeness of computer models could increase their performance in tasks at which humans excel (Sinz et al., 2019; Hassabis et al., 2017; Ullman, 2019; Toneva & Wehbe, 2019; Yang et al., 2022).

### Dataset limitations

A major limitation of our dataset is the backward and forward noise introduced by the very short (200ms) stimulus onset asynchronies (SOAs) of the RSVP paradigm (Intraub, 1981; Keysers et al., 2001; Grootswagers et al., 2019). The forward noise at a given EEG image trial comes from the ongoing neural activity of the previous trial, whereas the backward noise coming from the following trial starts from around 260ms after image onset, which corresponds to the SOA length plus the amount of time required for the visual information to travel from the retina to the visual cortex. Despite these noise sources, we showed that the visual responses are successfully predicted during the entire EEG epoch. We believe that averaging the EEG image conditions across several repetitions of measurements reduced the noise, and that the backward noise was further mitigated given that the neural processing required to detect and recognize object categories can be achieved in the first 150ms of vision (Thorpe et al., 1996; Rousselet et al., 2002). A second limitation concerns the ecological validity of the dataset. The stimuli images used consisted of objects presented at foveal vision with natural backgrounds that have little clutter. Furthermore, participants were asked to constantly gaze at a central fixation target. This does not truthfully represent human vision, in which objects are perceived and recognized also when at the periphery of the visual field, within cluttered scenes, and while making eye movements. However, our results pave the way towards studies aiming to provide large amounts of EEG responses recorded during more natural viewing conditions.

### Contribution to the THINGS initiative

The visual brain is an ensemble of billions of neurons communicating with high spatial and temporal precision. However, neither current neural recording modalities, nor single lab efforts can capture this complexity. This motivates the need to integrate data across different imaging modalities and labs. To address this challenge, the so-called THINGS initiative promotes using the THINGS database to collect and share behavioral and neuroscientific datasets for the same set of images - also used here - among vision researchers (https://things-initiative.org/). We contribute to the initiative by providing rich high temporal resolution EEG data, that complements other datasets in both a within- and between-modality fashion. As an example of the within-modality fashion, Grootswagers and collaborators recently published an EEG dataset of visual responses to images coming from the THINGS database (Grootswager et al., 2022). While their dataset comprises more participants and image conditions, our dataset provides more repetitions of measurements, longer image presentation latencies, and an extensive assessment of the dataset’s potential based on the resulting high signal-to-noise ratio. Researchers can choose between one or the other based on the nature, requirements and constraints of their own experiments. As an example of the between-modality fashion, our data can be used to make bridges from the EEG temporal domain to, for example, the fMRI spatial domain through modeling frameworks such as representational similarity analysis (Kriegeskorte et al., 2008; Cichy et al., 2014; Cichy et al., 2016; Khaligh-Razavi et al., 2017), thus promoting a more integrated understanding of the neural basis of visual object recognition.

### Comparing the modeling results of the four DNNs evaluated

The size and quality of our dataset make it a good candidate for the comparison of predictive and explanatory models of the visual brain (Schrimpf et al., 2020; Cichy et al., 2019). Here, we built encoding models using four DNNs: despite the prediction accuracies of these DNNs being overall qualitatively similar (Storrs et al., 2021), the results of our analyses suggest that the EEG data is best predicted by the linearizing encoding models based on the recurrent CORnet-S architecture (Kubilius et al., 2019). This supports a growing amount of literature which asserts that recurrent computations are critical for object recognition along the ventral stream, and therefore any model of visual object processing must also take recurrency into account (Kriegeskorte, 2015; Spoerer et al., 2017; Mohsenzadeh et al., 2018; Kar et al., 2019; Kietzmann et al., 2019b; Kubilius et al., 2019; Rajaei et al., 2019; van Bergen & Kriegeskorte, 2020). However, this interpretation should be taken with a grain of salt as we compared DNNs differing not only in the hypotheses of visual processing they embedded (e.g., recurrent vs. pure feedforward visual processing), but also in potential confounding factors such as architecture and complexity.

### The modeling accuracy is not homogeneous across time

As expected, the prediction accuracies of our encoding algorithms did not reach the noise ceiling level (**Supplementary Figures 1-2**), indicating that our dataset is well suited for further model improvements. Interestingly, we found that the modeling accuracy is not homogeneous across time: the differences between the prediction accuracy and the noise ceiling are smaller in the first 100ms after image onset, and peak at 200-220ms, suggesting that the four DNNs used are more similar to the brain at earlier stages of visual processing. This calls for future improvements in model building (e.g., by including high-level visual semantics or improving biological realism of the models) to more closely match the internal representations of the brain at all time points.

### Conclusion

We view our EEG dataset as a valuable tool for computational neuroscientists and computer scientists. We believe that its largeness, richness and quality will facilitate steps towards a deeper understanding of the neural mechanisms underlying visual processing and towards more human-like artificial intelligence models.

## Materials and methods

### Participants

Ten healthy adults (mean age 28.5 years, SD=4; 8 female, 2 male) participated, all having normal or corrected-to-normal vision. They all provided informed written consent and received monetary reimbursement. Procedures were approved by the ethical committee of the Department of Education and Psychology at Freie Universität Berlin and were in accordance with the Declaration of Helsinki.

### Stimuli

All images came from THINGS (Hebart et al., 2019), a database of 12 or more images of objects on a natural background for each of 1,854 object concepts, where each concept (e.g., antelope, strawberry, t-shirt) belongs to one of 27 higher-level categories (e.g., animal, food, clothing). The building of encoding models involves two stages: model training and model evaluation. Since each of these stages requires an independent data partition, we pseudo-randomly divided the 1,854 object concepts into non-overlapping 1,654 training and 200 test concepts under the constraint that the same proportion of the 27 higher-level categories had to be kept in both partitions. We then selected ten images for each training partition concept and one image for each test partition concept, resulting in a training image partition of 16,540 image conditions (1,654 training object concepts × 10 images per concept = 16,540 training image conditions) and a test image partition of 200 image conditions (200 test object concepts × 1 image per concept = 200 test image conditions). We used the training and test data partitions for the encoding model training and testing, respectively. The experiment had an orthogonal target detection task (see “experimental paradigm” section below), and as task-relevant target stimuli we used 10 different images of the “Toy Story” character Buzz Lightyear. All images were of square size. We reshaped them to 500 × 500 pixels for the EEG data collection paradigm. For the modeling with DNNs we reshaped the images to 224 × 224 pixels, and normalized them.

### Experimental Paradigm

The experiment consisted in a RSVP paradigm (Intraub, 1981; Keysers et al., 2001; Grootswagers et al., 2019) with an orthogonal target detection task to ensure participants paid attention to the visual stimuli. All 10 participants completed four equivalent experimental sessions, resulting in 10 datasets of 16,540 training images conditions repeated 4 times and 200 test image conditions repeated 80 times, for a total of (16,540 training image conditions × 4 training image repetitions) + (200 test image conditions × 80 test image repetitions) = 82,160 image trials per dataset.

One session comprised 19 runs, all lasting around 5m. In each of the first 4 runs we showed participants the 200 test image conditions through 51 rapid serial sequences of 20 images, for a total of 4 test runs × 51 sequences per run × 20 images per sequence = 4,080 image trials. In each of the following 15 runs we showed 8,270 training image conditions (half of all the training image conditions, as different halves were shown on different sessions) through 56 rapid serial sequences of 20 images, for a total of 15 training runs × 56 sequences per run × 20 images per sequence = 16,800 image trials.

Every rapid serial sequence started with 750ms of blank screen, then each of the 20 images was presented centrally with a visual angle of 7 degrees for 100ms and a SOA of 200ms, and it ended with another 750ms of blank screen. After every rapid sequence there were up to 2s during which we instructed participants to first blink (or make any other movement) and then report, with a keypress, whether the target image of Buzz Lightyear appeared in the sequence. The images were presented in a pseudo-randomized order, and a target image appeared in 6 sequences per run. A central bull’s eye fixation target (Thaler et al., 2013) was present on the screen throughout the entire experiment, and we asked participants to constantly gaze at it. We controlled stimulus presentation using the Psychtoolbox (Brainard, 1997), and recorded EEG data during the experimental sessions.

### EEG recording and preprocessing

We recorded the EEG data using a 64-channel EASYCAP with electrodes arranged in accordance with the standard 10-10 system (Nuwer et al., 2998), and a Brainvision actiCHamp amplifier. We recorded the data at a sampling rate of 1000Hz, while performing online filtering (between 0.1Hz and 100Hz) and referencing (to the Fz electrode). We performed offline preprocessing in Python, using the MNE package (Gramfort et al., 2013). We epoched the continuous EEG data into trials ranging from 200ms before stimulus onset to 800ms after stimulus onset, and applied baseline correction by subtracting the mean of the pre-stimulus interval for each trial and channel separately. We then down-sampled the epoched data to 100Hz, and we selected 17 channels overlying occipital and parietal cortex for further analysis (O1, Oz, O2, PO7, PO3, POz, PO4, PO8, P7, P5, P3, P1, Pz, P2, P4, P6, P8). All trials containing target stimuli were not analyzed further, and we randomly selected and retained 4 measurement repetitions for each training image condition and 80 measurement repetitions for each test image condition. Next, we applied multivariate noise normalization (Guggenmos et al., 2018) independently to the data of each recording session. For each participant, the preprocessing resulted in the EEG *biological training* (BioTrain) data matrix of shape (16,540 training image conditions × 4 condition repetitions × 17 EEG channels × 100 EEG time points) and *biological test* (BioTest) data matrix of shape (200 test image conditions × 80 condition repetitions × 17 EEG channels × 100 EEG time points). We used the BioTrain and BioTest EEG data for the encoding models training and testing, respectively.

### DNN models used

We built linearizing encoding models (Wu et al., 2006; Kay et al., 2008; Naselaris et al., 2011; van Gerven, 2017; Kriegeskorte & Douglas, 2019) of EEG visual responses using four different DNNs: AlexNet (Krizhevsky, 2014), a supervised feedforward neural network of 5 convolutional layers followed by 3 fully-connected layers that won the Imagenet large-scale visual recognition challenge in 2012; ResNet-50 (He et al, 2016), a supervised feedforward 50 layer neural network with shortcut connections between layers at different depths; CORnet-S (Kubilius et al., 2019), a supervised deep recurrent neural network of four convolutional layers and one fully-connected layer; MoCo (Chen et al., 2020), a feedforward ResNet-50 architecture trained in a self-supervised fashion. All of them had been pre-trained on object categorization on the ILSVRC-2012 training image partition (Russakovsky et al., 2015).

### Linearizing encoding models of EEG visual responses

The first step in building linearizing encoding models is to use DNNs to non-linearly transform the image input space onto a feature space. A DNNs feature space is given by its feature maps, layerwise representations (non-linear transformations) of the input images. To get the training and test feature maps we fed the training and test images separately to each DNN and appended the vectorized image representations of its layers onto each other. We extracted AlexNet’s feature maps from layers maxpool1, maxpool2, ReLU3, ReLU4, maxpool5, ReLU6, ReLU7, and fc8; ResNet-50’s and MoCo’s feature maps from the last layer of each of their four blocks, and from the decoder layer; CORnet-S’ feature maps from the last layers of areas V1, V2 (at both time points), V4 (at all four time points), IT (at both time points), and from the decoder layer. We then standardized the appended feature maps of the training and test data to zero mean and unit variance for each feature across the sample (images) dimension, using the mean and standard deviation of the training feature maps. Finally, we used the Scikit-learn (Pedregosa et al., 2011) implementation of non-linear principal component analysis (computed on the training feature maps using a polynomial kernel of degree 4) to reduce the feature maps of both the training and test images to 1,000 components. For each DNN model, this resulted in the training feature maps matrix of shape (16,540 training image conditions × 1,000 features) and test feature maps matrix of shape (200 test image conditions × 1,000 features).

The second step in building linearizing encoding models is to linearly map the DNNs’ feature space onto the EEG neural space, effectively predicting the EEG responses to images. We performed this linear mapping independently for each participant, DNN model and EEG feature (i.e., for each of the 17 EEG channels (**c**) × 100 EEG time points (**t**) = 1,700 EEG features). We fitted the weights ***W***_***t,c***_ of a linear regression using the DNNs’ training feature maps as the predictors and the corresponding BioTrain data (averaged across the image conditions repetitions) as the criterion: during training the regression weights learned the existing linear relationship between the DNN feature maps of a given image and the EEG responses of that same image. No regularization techniques were used. We multiplied ***W***_***t,c***_ with the DNNs’ test feature maps. For each participant and DNN, this resulted in the *synthetic test* (SynTest) EEG data matrix of shape (200 test image conditions × 17 EEG channels × 100 EEG time points).

### Correlation

We used a Pearson correlation to assess how similar the SynTest EEG data of each participant and DNN is to the corresponding BioTest data, thus quantifying the encoding models’ predicted power. We started the analysis by averaging the BioTest data across 40 image conditions repetitions (see “noise ceiling” section below), resulting in a BioTest data matrix equivalent in shape to the SynTest data matrix (200 test image conditions × 17 EEG channels × 100 EEG time points). Next, we implemented a nested loop over the EEG channels and time points. At each loop iteration we indexed the 200-dimensional BioTest data vector containing the 200 test image conditions of the EEG channel (**c**) and time point (**t**) in question, and correlated it with the corresponding 200-dimensional SynTest data vector. This procedure yielded a Pearson correlation coefficient matrix of shape (17 EEG channels × 100 EEG time points). Finally, we averaged the Pearson correlation coefficient matrix over the EEG channels, obtaining a correlation results vector of length (100 EEG time points) for each participant and DNN.

### Pairwise decoding

The rationale of this analysis was to see if a classifier trained on the BioTest data is capable of generalizing its performance to the SynTest data. This is a complementary way (to the correlation analysis) to assess the similarity between the SynTest data and the BioTest data, hence the encoding models’ predictive power. We started the analysis by averaging 40 BioTest data image conditions repetitions (see “noise ceiling” section below) into 10 pseudo-trials of 4 repeats each, yielding a matrix of shape (200 test image conditions × 10 image condition pseudo-trials × 17 EEG channels × 100 EEG time points). Next, we used the pseudo-trials for training linear SVMs to perform binary classification between each pair of the 200 BioTest data image conditions (for a total of 19,900 image condition pairs) using their EEG channels vectors (of 17 components). We then tested the trained classifiers on the corresponding pairs of SynTest data image conditions. We performed the pairwise decoding analysis independently for each EEG time point (**t**), which resulted in a matrix of decoding accuracy scores of shape (19,900 image condition pairs × 100 EEG time points). We then averaged the decoding accuracy scores matrix across the image condition pairs, obtaining a pairwise decoding results vector of length (100 EEG time points) for each participant and DNN.

### Zero-shot identification

In this analysis we exploited the linearizing encoding models’ predictive power to identify the BioTest data image conditions in a zero-shot fashion, that its, to identify arbitrary image conditions without prior training (Kay et al., 2008; Seeliger et al., 2017; Horikawa & Kamitani, 2017). We identified each BioTest data image condition using the SynTest data and an additional synthesized EEG dataset of up to 150,000 candidate image conditions. These 150,000 image conditions came from the ILSVRC-2012 (Russakovsky et al., 2015) validation (50,000) plus test (100,000) sets. We synthesized them into their corresponding EEG responses following the same procedure described above, resulting in the *synthetic Imagenet* (SynImagenet) data matrix of shape (150,000 image conditions × 17 EEG channels × 100 EEG time points). The zero-shot identification analysis involved two steps: feature selection and identification.

In the feature selection step we used the training data to pick only the most relevant EEG features (out of all 17 EEG channels × 100 EEG time points = 1,700 EEG features). We synthesized the EEG responses to the 16,540 training images, obtaining the *synthetic train* (SynTrain) data matrix of shape (16,540 training image conditions × 17 EEG channels × 100 EEG time points). Next, we correlated each SynTrain data feature (across the 16,540 training image conditions, with a Pearson correlation), with the corresponding BioTrain data feature (averaged across the image conditions repetitions). We then selected only the 300 BioTest data, SynTest data and SynImagenet data EEG features corresponding to the 300 highest correlation scores, thus obtaining a BioTest data matrix of shape (200 test image conditions × 80 condition repetitions × 300 EEG features), a SynTest data matrix of shape (200 test image conditions × 300 EEG features), and a SynImagenet data matrix of shape (150,000 image conditions × 300 EEG features).

In the identification step we started by averaging the BioTest data across all the 80 image conditions repetitions: this yielded feature vectors of 300 components for each of the 200 image conditions. Next, we correlated (through a Pearson correlation) the feature vectors of each BioTest data image condition with the feature vectors of all the candidate image conditions: the SynTest data image conditions plus a varying amount of SynImagenet data image conditions. We increased the set sizes of the SynImagenet candidate image conditions from 0 to 150,000 with steps of 1,000 images (for a total of 151 set sizes), where 0 corresponded to using only the SynTest data candidate image conditions, and performed the zero-shot identification at every set size. At each SynImagenet set size a BioTest data image condition is considered correctly identified if the correlation coefficient between its channel vector and the channel vector of the corresponding SynTest data image condition is higher than the correlation coefficients between its channel vector and the channel vectors of all other candidate SynTest data and SynImagenet data image conditions. Thus, we calculated the zero-shot identification accuracies through the ratio of correctly classified images over all 200 BioTest images, obtaining a zero-shot identification results vector of length (151 candidate image set sizes). We iterated the identification step 100 times, while always randomly selecting different SynImagenet data image conditions at each set size, and then averaged the results across the 100 iterations.

To extrapolate the drop in identification accuracy with larger candidate image set sizes we fit the power-law function to the results of each participant. The power law function is defined as:

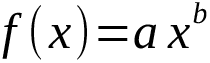

where *x* is the image set size, *a* and *b* are constants learned during function fitting, and *f* (*x*) is the predicted zero-shot identification accuracy. We fit the function using the 100 SynImagenet set sizes ranging from 50,200 to 150,200 images (along with their corresponding identification accuracies), and then used it to extrapolate the image set size required for the identification accuracy to drop below 10% and 0.5%.

### End-to-end encoding models of EEG visual responses

We based our end-to-end encoding models (Seeliger et al., 2021; Khosla et al., 2021; Allen et al., 2021) on the AlexNet architecture which, once trained, predicted the EEG responses to the test images. To match AlexNet’s output with the channel responses of our EEG data, we replaced AlexNet’s 1000-neurons output layer with a 17-neurons layer, where each neuron represented one of the 17 EEG channels. Next, we randomly initialized independent AlexNet instances for each participant and EEG time point (t). We used Pytorch (Paszke et al., 2019) to train the AlexNets on a regression task: given the input training images and the corresponding target BioTrain EEG data channel activity (averaged across the image condition repetitions), the models had to optimize their weights so to minimize the summed squared error between their predictions and the BioTrain data. We trained the models using batch sizes of 256 images and an Adam optimizer with a learning rate of 0.0001, a weight decay term of 0.001, and the default value for the remaining parameters. We implemented a cross-validation loop over the 200 test image conditions to identify the optimal amount of training epochs for the synthesis of each image’s EEG responses. At every loop iteration we selected the image condition of interest, synthesized the EEG responses to the remaining 199 test images for each of 30 training epochs, and correlated the synthetic data with the corresponding 199 biological test EEG data conditions, resulting in one correlation score per epoch. We then synthesized the EEG responses to the image condition of interest using the model weights of the epoch leading to the highest correlation score. For each participant, this resulted in the SynTest data matrix of shape (200 test image conditions × 17 EEG channels × 100 EEG time points).

### Noise ceiling calculation

We calculated the noise ceilings of the correlation and pairwise decoding analyses to estimate the theoretical maximum results given the level of noise in the BioTest data. If the results of the SynTest data reach this theoretical maximum the encoding models are successful in explaining all the BioTest data variance which can be explained. If not, further model improvements could lead to more accurate predictions of neural data.

For the noise ceiling estimation we randomly divided the BioTest data into two non-overlapping partitions of 40 image condition repetitions each, where the first partition corresponded to the 40 repeats of BioTest data image conditions used in the correlation and pairwise decoding analyses described above. We then performed the two analyses while substituting the SynTest data with the second BioTest data partition (averaged across image condition repetitions). This resulted in the noise ceiling lower bound estimates. To calculate the upper bound estimates we substituted the SynTest data with the average of the BioTest data over all 80 image condition repetitions and reiterated the two analyses. We assume that the true noise ceiling is somewhere in between the lower and the upper bound estimates. To avoid the results being biased by one specific configuration of the BioTest data repeats we iterated the correlation and pairwise decoding analyses 100 times, while always selecting different repeats for the two BioTest data partitions, and then averaged the results across the 100 iterations.

### Statistical testing

To assess the statistical significance of the correlation, pairwise decoding and zero-shot identification analyses we tested all results against chance using one-sample one-sided t-tests. Here, the rationale was to reject the null hypothesis H0 of the analyses results being at chance level with a confidence of 95% or higher (i.e., with a *P*-value of *P* < 0.05), thus supporting the experimental hypothesis H1 of the results being significantly higher than chance. The chance level differed across analyses: 0 in the correlation; 50% in the pairwise decoding; (1 / (200 test image conditions + *N* ILSVRC-2012 image conditions)) in the zero-shot identification (where *N* varied from 0 to 150,000). When analyzing the linearizing encoding models’ prediction accuracy using different amounts of training data we used a two-way repeated measures ANOVA to reject the null hypothesis H0 of no significant effects of number of image conditions and/or condition repetitions on the prediction accuracy, and a repeated measures two-sided t-test to reject the null hypothesis H0 of no significant differences between the effects of training image conditions and condition repetitions.

We controlled familywise error rate by applying a conservative Bonferroni-correction to the resulting *P*-values to correct for the number of EEG time points (*N* = 100) in the correlation and pairwise decoding analyses, for the amount of training data quartiles (*N* = 4) in the analysis of the linearizing encoding models’ prediction accuracy as a function of training image conditions and condition repetitions, and for the number of candidate images set sizes (*N* = 151) in the zero-shot identification analysis.

To calculate the confidence intervals of each statistic, we created 10,000 bootstrapped samples by sampling the participant-specific results with replacement. This yielded empirical distributions of the results, from which we took the 95% confidence intervals.

## Acknowledgments

A.T.G. is supported by a PhD fellowship of the Einstein Center for Neurosciences. G.R. is supported by the Alfons and Gertrud Kassel Foundation. R.M.C. is supported by German Research Council (DFG) grants (CI 241/1-1, CI 241/3-1, CI 241/1-7) and the European Research Council (ERC) starting grant (ERC-StG-2018-803370). We thank Martin Hebart for support with the THINGS database. We thank Daniel Kaiser and Kendrick Kay for helpful comments on the manuscript. We thank the HPC Service of ZEDAT, Freie Universität Berlin, for computing time.

## Author contributions

A.T.G., K.D. and R.M.C. designed research, A.T.G. acquired data, A.T.G. analyzed data, A.T.G., K.D., G.R. and R.M.C. interpreted results, A.T.G. prepared figures, A.T.G. drafted manuscript, A.T.G., K.D., G.R. and R.M.C. edited and revised manuscript. All authors approved the final version of the manuscript.

## Data availability

The raw and preprocessed EEG dataset, the stimuli image set and the extracted DNN feature maps are available on OSF.

## Code availability

The code to reproduce all the results is available on GitHub.

## References

Abiri R, Borhani S, Sellers EW, Jiang Y, Zhao X. 2019. A comprehensive review of EEG-based brain– computer interface paradigms. Journal of Neural Engineering, 16(1):011001. DOI: https://doi.org/10.1088/1741-2552/aaf12e

Allen EJ, St-Yves G, Wu Y, Breedlove JL, Prince JS, Dowdle LT, Nau M, Caron B, Pestilli F, Charest I, Hutchinson JB, Naselaris T, Kay K. 2022. A massive 7T fMRI dataset to bridge cognitive neuroscience and computational intelligence. Nature Neuroscience, 25(1):116–126. DOI: https://doi.org/10.1038/s41593-021-00962-x

Bankson BB, Hebart MN, Groen II, Baker CI. 2018. The temporal evolution of conceptual object representations revealed through models of behavior, semantics and deep neural networks. NeuroImage, 178:172–182. DOI: https://doi.org/10.1016/j.neuroimage.2018.05.037

Brainard DH. 1997. The psychophysics toolbox. Spatial Vision, 10:433–436. DOI: https://doi.org/10.1163/156856897X00357

Cadieu CF, Hong H, Yamins DLK, Pinto N, Ardila D, Solomon EA, Majaj NJ, DiCarlo JJ. 2014. Deep neural networks rival the representation of primate IT cortex for core visual object recognition. PLoS Computational Biology, 10(12):e1003963. DOI: https://doi.org/10.1371/journal.pcbi.1003963

Carandini M, Demb JB, Mante V, Tolhurst DJ, Dan Y, Olshausen BA, Gallant JL, Rust NC. 2005. Do we know what the early visual system does?. Journal of Neuroscience, 25(46):10577–10597. DOI: https://doi.org/10.1523/JNEUROSCI.3726-05.2005

Chang N, Pyles JA, Marcus A, Gupta A, Tarr M, Aminoff EM. 2019. BOLD5000, a public fMRI dataset while viewing 5000 visual images. Scientific Data, 6(1):1–18. DOI: https://doi.org/10.1038/s41597-019-0052-3

Charest I, Kievit RA, Schmitz TW, Deca D, Kriegeskorte N. 2014. Unique semantic space in the brain of each beholder predicts perceived similarity. Proceedings of the National Academy of Sciences, 111(40): 14565–14570. DOI: https://doi.org/10.1073/pnas.1402594111

Chen X, Fan H, Girshick R, He K. 2020. Improved baselines with momentum contrastive learning. arXiv preprint, arXiv:2003.04297. DOI: https://doi.org/10.48550/arXiv.2003.04297

Cichy RM, Kaiser D. 2019. Deep neural networks as scientific models. Trends in Cognitive Sciences, 23(4):305–317. DOI: https://doi.org/10.1016/j.tics.2019.01.009

Cichy RM, Khosla A, Pantazis D, Torralba A, Oliva A. 2016. Comparison of deep neural networks to spatio-temporal cortical dynamics of human visual object recognition reveals hierarchical correspondence. Scientific Reports, 6(1):1–13. DOI: https://doi.org/10.1038/srep27755

Cichy RM, Pantazis D, Oliva A. 2014. Resolving human object recognition in space and time. Nature Neuroscience, 17(3):455–462. DOI: https://doi.org/10.1038/nn.3635

Cichy RM, Roig G, Oliva A. 2019. The Algonauts Project. Nature Machine Intelligence, 1(12):613–613. DOI: https://doi.org/10.1038/s42256-019-0127-z

Dijkstra N, Mostert P, de Lange FP, Bosch S, Gerven MA. 2018. Differential temporal dynamics during visual imagery and perception. Elife, 7:e33904. DOI: https://doi.org/10.7554/eLife.33904

Fukushima K, Miyake S. 1982. Neocognitron: A self-organizing neural network model for a mechanism of visual pattern recognition. In Competition and Cooperation in Neural Nets: 267–285. Springer, Berlin, Heidelberg. DOI: https://doi.org/10.1007/978-3-642-46466-9_18

Goodale MA, Milner AD. 1992. Separate visual pathways for perception and action. Trends in Neurosciences, 15(1): 20–25. DOI: https://doi.org/10.1016/0166-2236(92)90344-8

Gramfort A, Luessi M, Larson E, Engemann DA, Strohmeier D, Brodbeck C, Goj R, Jas M, Brooks T, Parkkonen L, Hämäläinen MS. 2013. MEG and EEG data analysis with MNE-Python. Frontiers in Neuroscience, 7(267):1–13. DOI: https://doi.org/10.3389/fnins.2013.00267

Grill-Spector K, Kourtzi Z, Kanwisher N. 2001. The lateral occipital complex and its role in object recognition. Vision Research, 41(10-11):1409–1422. DOI: https://doi.org/10.1016/S0042-6989(01)00073-6

Grootswagers T, Robinson AK, Carlson TA. 2019. The representational dynamics of visual objects in rapid serial visual processing streams. NeuroImage, 188:668–679. DOI: https://doi.org/10.1016/j.neuroimage.2018.12.046

Grootswagers T, Zhou I, Robinson AK, Hebart MN, Carlson TA. 2022. Human EEG recordings for 1,854 concepts presented in rapid serial visual presentation streams. Scientific Data, 9(1):1–7. DOI: https://doi.org/10.1038/s41597-021-01102-7

Güçlü U, van Gerven MAJ. 2015. Deep neural networks reveal a gradient in the complexity of neural representations across the ventral stream. Journal of Neuroscience, 35(27):10005–10014. DOI: https://doi.org/10.1523/JNEUROSCI.5023-14.2015

Guest O, Martin AE. 2021. On logical inference over brains, behaviour, and artificial neural networks.

Guggenmos M, Sterzer P, Cichy RM. 2018. Multivariate pattern analysis for MEG: A comparison of https://doi.org/10.1016/j.neuroimage.2018.02.044

Harel A, Groen II, Kravitz DJ, Deouell LY, Baker CI. 2016. The temporal dynamics of scene https://doi.org/10.1523/ENEURO.0139-16.2016

Hassabis D, Kumaran D, Summerfield C, Botvinick M. 2017. Neuroscience-inspired artificial intelligence. Neuron, 95(2):245–258. DOI: https://doi.org/10.1016/j.neuron.2017.06.011

Haxby JV, Guntupalli JS, Nastase SA, Feilong M. 2020. Hyperalignment: Modeling shared information encoded in idiosyncratic cortical topographies. Elife, 9:e56601. DOI: https://doi.org/10.7554/eLife.56601

Haynes JD, Rees G. 2006. Decoding mental states from brain activity in humans. Nature Reviews Neuroscience, 7(7):523–534. DOI: https://doi.org/10.1038/nrn1931

He K, Zhang X, Ren S, Sun J. 2016. Deep residual learning for image recognition. Proceedings of the IEEE Conference on Computer Vision and Pattern Recognition, 770–778. DOI: https://doi.org/10.1109/CVPR.2016.90

Hebart MN, Dickter AH, Kidder A, Kwok WY, Corriveau A, Van Wicklin C, Baker CI. 2019. THINGS: A database of 1,854 object concepts and more than 26,000 naturalistic object images. PLoS ONE, 14(10): e0223792. DOI: https://doi.org/10.1371/journal.pone.0223792

Horikawa T, Kamitani Y. 2017. Generic decoding of seen and imagined objects using hierarchical visual features. Nature Communications, 8(1):1–15. DOI: https://doi.org/10.1038/ncomms15037

Intraub H. 1981. Rapid conceptual identification of sequentially presented pictures. Journal of Experimental Psychology: Human Perception and Performance, 7(3):604. DOI: https://doi.org/10.1037/0096-1523.7.3.604

Kar K, Kubilius J, Schmidt K, Issa EB, DiCarlo JJ. 2019. Evidence that recurrent circuits are critical to the ventral stream’s execution of core object recognition behavior. Nature Neuroscience, 22(6):974–983. DOI: https://doi.org/10.1038/s41593-019-0392-5

Kay KN, Naselaris T, Prenger RJ, Gallant JL. 2008. Identifying natural images from human brain activity. Nature, 452(7185):352–355. DOI: https://doi.org/10.1038/nature06713

Keysers C, Xiao DK, Földiák P, Perrett DI. 2001. The speed of sight. Journal of cognitive neuroscience, 13(1):90–101. DOI: https://doi.org/10.1162/089892901564199

Khaligh-Razavi SM, Henriksson L, Kay K, Kriegeskorte N. 2017. Fixed versus mixed RSA: Explaining visual representations by fixed and mixed feature sets from shallow and deep computational https://doi.org/10.1016/j.jmp.2016.10.007

Khosla M, Ngo GH, Jamison K, Kuceyeski A, Sabuncu MR. 2021. Cortical response to naturalistic stimuli is largely predictable with deep neural networks. Science Advances, 7(22):eabe7547. DOI: https://doi.org/10.1126/sciadv.abe7547

Kietzmann TC, McClure P, Kriegeskorte N. 2019a. Deep neural networks in computational neuroscience. In Oxford Research Encyclopedia of Neuroscience. DOI: https://doi.org/10.1093/acrefore/9780190264086.013.46

Kietzmann TC, Spoerer CJ, Sörensen LK, Cichy RM, Hauk O, Kriegeskorte N. 2019b. Recurrence is required to capture the representational dynamics of the human visual system. Proceedings of the National Academy of Sciences, 116(43):21854–21863. DOI: https://doi.org/10.1073/pnas.1905544116

Kriegeskorte N. 2015. Deep neural networks: a new framework for modeling biological vision and brain information processing. Annual Review of Vision Science, 1:417–446. DOI: https://doi.org/10.1146/annurev-vision-082114-035447

Kriegeskorte N, Douglas PK. 2019. Interpreting encoding and decoding models. Current opinion in neurobiology, 55, 167–179. DOI: https://doi.org/10.1016/j.conb.2019.04.002

Kriegeskorte N, Mur M, Bandettini PA. 2008. Representational similarity analysis-connecting the branches of systems neuroscience. Frontiers in Systems Neuroscience, 2:4.8 DOI: https://doi.org/10.3389/neuro.06.004.2008

Krizhevsky A. 2014. One weird trick for parallelizing convolutional neural networks. arXiv preprint, arXiv:1404.5997

Kubilius J, Schrimpf M, Kar K, Rajalingham R, Hong H, Majaj N, Issa E, Bashivan P, Prescott-Roy J, Schmidt K, Nayebi A, Bear D, Yamins DL, DiCarlo JJ. 2019. Brain-like object recognition with high-performing shallow recurrent ANNs. Advances in neural information processing systems, 32.

Kwon OY, Lee MH, Guan C, Lee SW. 2019. Subject-independent brain–computer interfaces based on deep convolutional neural networks. IEEE transactions on neural networks and learning systems, 31(10):3839–3852. DOI: https://doi.org/10.1109/TNNLS.2019.2946869

Logothetis NK, Sheinberg DL. 1996. Visual object recognition. Annual review of neuroscience, 19(1):577–621. DOI: https://doi.org/10.1146/annurev.ne.19.030196.003045

Malach R, Levy I, Hasson U. 2002. The topography of high-order human object areas. Trends in Cognitive Sciences, 6(4):176–184. DOI: https://doi.org/10.1016/S1364-6613(02)01870-3

Marr D. 1980. Visual information processing: The structure and creation of visual representations. Philosophical Transactions of the Royal Society of London. B, Biological Sciences, 290(1038):199–218. DOI: https://doi.org/10.1098/rstb.1980.0091

Mohsenzadeh Y, Qin S, Cichy RM, Pantazis D. 2018. Ultra-Rapid serial visual presentation reveals dynamics of feedforward and feedback processes in the ventral visual pathway. Elife, 7:e36329. DOI: https://doi.org/10.7554/eLife.36329

Mur M, Bandettini PA, Kriegeskorte N. 2009. Revealing representational content with pattern-information fMRI—an introductory guide. Social Cognitive and Affective Neuroscience, 4(1):101–109. DOI: https://doi.org/10.1093/scan/nsn044

Naselaris T, Allen E, Kay K. 2021. Extensive sampling for complete models of individual brains. DOI: https://doi.org/10.1016/j.cobeha.2020.12.008

Naselaris T, Kay KN, Nishimoto S, Gallant JL. 2011. Encoding and decoding in fMRI. NeuroImage, 56(2):400–410. DOI: https://doi.org/10.1016/j.neuroimage.2010.07.073

Naselaris T, Olman CA, Stansbury DE, Ugurbil K, Gallant JL. 2015. A voxel-wise encoding model for early visual areas decodes mental images of remembered scenes. NeuroImage, 105:215–228. DOI: https://doi.org/10.1016/j.neuroimage.2014.10.018

Nuwer MR, Comi G, Emerson R, Fuglsang-Frederiksen A, Guérit JM, Hinrichs H, Ikeda A, Luccas FJC, Rappelsburger P. 1998. IFCN standards for digital recording of clinical EEG. Electroencephalography and Clinical Neurophysiology, 106(3):259–261. DOI: https://doi.org/10.1016/s0013-4694(97)00106-5

Paszke A, Gross S, Massa F, Lerer A, Bradbury J, Chanan G, Killeen T, Lin Z, Gimelshein N, Antiga L, Desmaison A. 2019. Pytorch: An imperative style, high-performance deep learning library. Advances in Neural Information Processing Systems, 32:8026–8037.

Pedregosa F, Varoquaux G, Gramfort A, Michel V, Thirion B, Grisel O, Blondel M, Prettenhofer P, Weiss R, Dubourg V, Vanderplas J, Passos A, Cournapeau D, Brucher M, Perrot M, Duchesnay É. 2011. Scikit-learn: Machine learning in Python. the Journal of Machine Learning Research, 12:2825–2830.

Petit J, Rouillard J, Cabestaing F. 2021. EEG-based brain–computer interfaces exploiting steady-state somatosensory-evoked potentials: a literature review. Journal of Neural Engineering, 18(5):051003. DOI: https://doi.org/10.1088/1741-2552/ac2fc4

Rajaei K, Mohsenzadeh Y, Ebrahimpour R, Khaligh-Razavi SM. 2019. Beyond core object recognition: Recurrent processes account for object recognition under occlusion. PLoS computational biology, 15(5):e1007001. DOI: https://doi.org/10.1371/journal.pcbi.1007001

Richard H, Gresele L, Hyvarinen A, Thirion B, Gramfort A, Ablin P. 2020. Modeling shared responses in neuroimaging studies through multiview ica. Advances in Neural Information Processing Systems, 33:19149–19162.

Richards BA, Lillicrap TP, Beaudoin P, Bengio Y, Bogacz R, Christensen A, Clopath C, Costa RP, de Berker A, Ganguli S, Gillon CJ. 2019. A deep learning framework for neuroscience. Nature Neuroscience, 22(11):1761–1770. DOI: https://doi.org/10.1038/s41593-019-0520-2

Riesenhuber M, Poggio T. 1999. Hierarchical models of object recognition in cortex. Nature neuroscience, 2(11):1019–1025. DOI: https://doi.org/10.1038/14819

Rousselet GA, Fabre-Thorpe M, Thorpe SJ. 2002. Parallel processing in high-level categorization of natural images. Nature Neuroscience, 5(7):629–630. DOI: https://doi.org/10.1038/nn866

Russakovsky O, Deng J, Su H, Krause J, Satheesh S, Ma S, Huang Z, Karpathy A, Khosla A, Bernstein M, Berg AC, Fei-Fei L. 2015. ImageNet Large Scale Visual Recognition Challenge. International Journal of Computer Vision, 115(3):211–252. DOI: https://doi.org/10.1007/s11263-015-0816-y

Saxe A, Nelli S, Summerfield C. 2021. If deep learning is the answer, what is the question?. Nature Reviews Neuroscience, 22(1):55–67. DOI: https://doi.org/10.1038/s41583-020-00395-8

Schrimpf M, Kubilius J, Lee MJ, Murty NAR, Ajemian R, DiCarlo JJ. 2020. Integrative benchmarking to advance neurally mechanistic models of human intelligence. Neuron, 108(3):413–423. DOI: https://doi.org/10.1016/j.neuron.2020.07.040

Seeliger K, Ambrogioni L, Güçlütürk Y, van den Bulk LM, Güçlü U, van Gerven MAJ. 2021. End-to-end neural system identification with neural information flow. PLOS Computational Biology, 17(2):e1008558. DOI: https://doi.org/10.1371/journal.pcbi.1008558

Seeliger K, Fritsche M, Güçlü U, Schoenmakers S, Schoffelen J-M, Bosch S, van Gerven, MAJ. 2017. Convolutional neural network-based encoding and decoding of visual object recognition in space and time. NeuroImage, 180:253–266. DOI: https://doi.org/10.1016/j.neuroimage.2017.07.018

Sinz FH, Pitkow X, Reimer J, Bethge M, Tolias AS. 2019. Engineering a less artificial intelligence. Neuron, 103(6):967–979. DOI: https://doi.org/10.1016/j.neuron.2019.08.034

Spoerer CJ, McClure P, Kriegeskorte N. 2017. Recurrent convolutional neural networks: a better model of biological object recognition. Frontiers in psychology, 8:1551. DOI: https://doi.org/10.3389/fpsyg.2017.01551

Storrs KR, Kietzmann TC, Walther A, Mehrer J, Kriegeskorte N. 2021. Diverse Deep Neural Networks All Predict Human Inferior Temporal Cortex Well, After Training and Fitting. Journal of Cognitive Neuroscience, 33(10):2044–2064. DOI: https://doi.org/10.1162/jocn_a_01755

Tanaka K. 1996. Inferotemporal cortex and object vision. Annual review of neuroscience, 19:109–139. DOI: https://doi.org/10.1146/annurev.ne.19.030196.000545

Thaler L, Schütz AC, Goodale MA, Gegenfurtner KR. 2013. What is the best fixation target? The effect of target shape on stability of fixational eye movements. Vision Research, 76:31–42. DOI: https://doi.org/10.1016/j.visres.2012.10.012

Thorpe S, Fize D, Marlot C. 1996. Speed of processing in the human visual system. Nature, 381(6582):520–522. DOI: https://doi.org/10.1038/381520a0

Toneva M, Wehbe L. 2019. Interpreting and improving natural-language processing (in machines) with natural language-processing (in the brain). Advances in Neural Information Processing Systems, 32.

Ullman S. 2000. High-level vision: Object recognition and visual cognition. MIT press.

Ullman S. 2019. Using neuroscience to develop artificial intelligence. Science, 363(6428):692–693. DOI: https://doi.org/10.1126/science.aau6595

Van Essen, D.C., Anderson, C.H. and Felleman, D.J., 1992. Information processing in the primate visual system: an integrated systems perspective. Science, 255(5043):419–423. DOI: https://doi.org/10.1126/science.1734518

van Bergen RS, Kriegeskorte N. 2020. Going in circles is the way forward: the role of recurrence in visual inference. Current Opinion in Neurobiology, 65:176–193. DOI: https://doi.org/10.1016/j.conb.2020.11.009

van de Nieuwenhuijzen ME, Backus AR, Bahramisharif A, Doeller CF, Jensen O, van Gerven MA. 2013. MEG-based decoding of the spatiotemporal dynamics of visual category perception. Neuroimage, 83:1063–1073. DOI: https://doi.org/10.1016/j.neuroimage.2013.07.075

van Gerven MA. 2017. A primer on encoding models in sensory neuroscience. Journal of Mathematical Psychology, 76:172–183. DOI: https://doi.org/10.1016/j.jmp.2016.06.009

Wu MC-K, David SV, Gallant JL. 2006. Complete functional characterization of sensory neurons by system identification. Annual Review of Neuroscience, 29(1):477–505. DOI: https://doi.org/10.1146/annurev.neuro.29.051605.113024

Yamins DLK, DiCarlo JJ. 2016. Using goal-driven deep learning models to understand sensory cortex. Nature Neuroscience, 19(3):356–365. DOI: https://doi.org/10.1038/nn.4244

Yamins DLK, Hong H, Cadieu CF, Solomon EA, Seibert D, DiCarlo JJ. 2014. Performance-optimized hierarchical models predict neural responses in higher visual cortex. Proceedings of the National Academy of Sciences, 111(23):8619–8624. DOI: https://doi.org/10.1073/pnas.1403112111

Yang X, Yan J, Wang W, Li S, Hu B, Lin J. 2022. Brain-inspired models for visual object recognition: an overview. Artificial Intelligence Review, 1–49. DOI: https://doi.org/10.1007/s10462-021-10130-z

Zhang K, Robinson N, Lee SW, Guan C. 2021. Adaptive transfer learning for EEG motor imagery classification with deep Convolutional Neural Network. Neural Networks, 136:1–10. DOI: https://doi.org/10.1016/j.neunet.2020.12.013

