## Supplementary material for "A large and rich EEG dataset for modeling human visual object recognition"

### SUPPLEMENTARY FIGURES

**Supplementary Figure 1:** Difference between the correlation results and the noise ceiling lower bound estimates ……..2

**Supplementary Figure 2:** Difference between the pairwise decoding results and the noise ceiling lower bound estimates ……..3

**Supplementary Figure 3:** Zero-shot identification, three most correlated candidate image conditions ……..4

**Supplementary Figure 4:** Zero-shot identification, ten most correlated candidate image conditions ……..5

**Supplementary Figure 5:** Effect of image conditions and condition repetitions on linearizing encoding models' prediction accuracy, individual participants' results……..6

**Supplementary Figure 6:** Contribution of image conditions and condition repetitions on linearizing encoding models' prediction accuracy, individual participants' results……..7

**Supplementary Figure 7:** Evaluating the prediction accuracy of linearizing encoding models which generalize to novel participants through correlation, individual participants' results……..8

**Supplementary Figure 8:** Evaluating the prediction accuracy of linearizing encoding models which generalize to novel participants through pairwise decoding, individual participants' results ……..9

**Supplementary Figure 9:** Evaluating the end-to-end encoding models' prediction accuracy through correlation, individual participants' results ……10

**Supplementary Figure 10:** Evaluating the end-to-end encoding models' prediction accuracy through pairwise decoding, individual participants' results ……..11

### SUPPLEMENTARY TABLES

**Supplementary Table 1:** Extrapolating the zero-shot identification accuracy, three most correlated candidate image conditions ……12

**Supplementary Table 2:** Extrapolating the zero-shot identification accuracy, ten most correlated candidate image conditions ……13

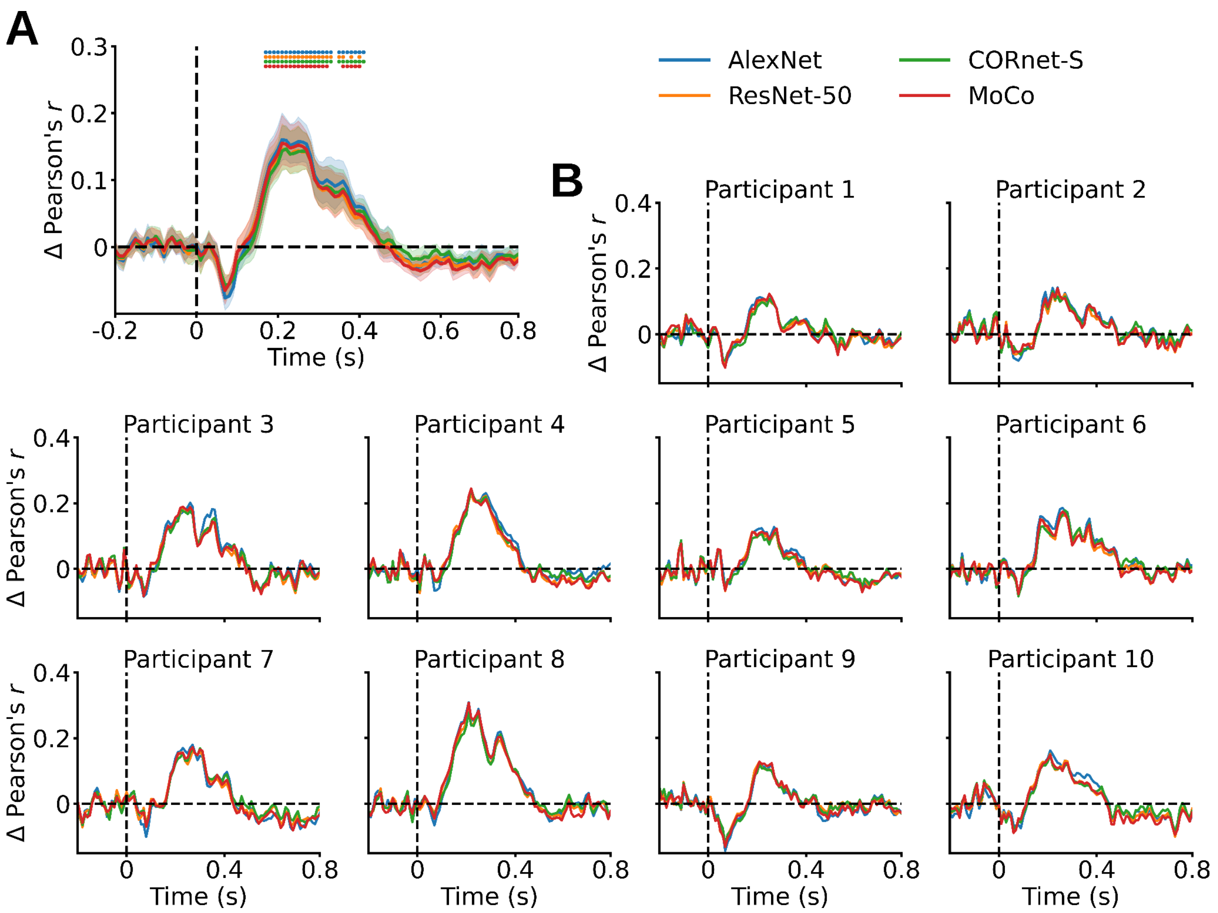


**Supplementary Figure 1.** Difference between the correlation results and the noise ceiling lower bound estimates. (**A**) The differences averaged across participants are significantly above zero between 170ms and 410ms, with peaks at 210-220ms ($P < 0.05$, one-sample one-sided t-test, Bonferroni-corrected). (**B**) Individual participants' results. Error margins, asterisks and black dashed lines as in **Figure 3**.

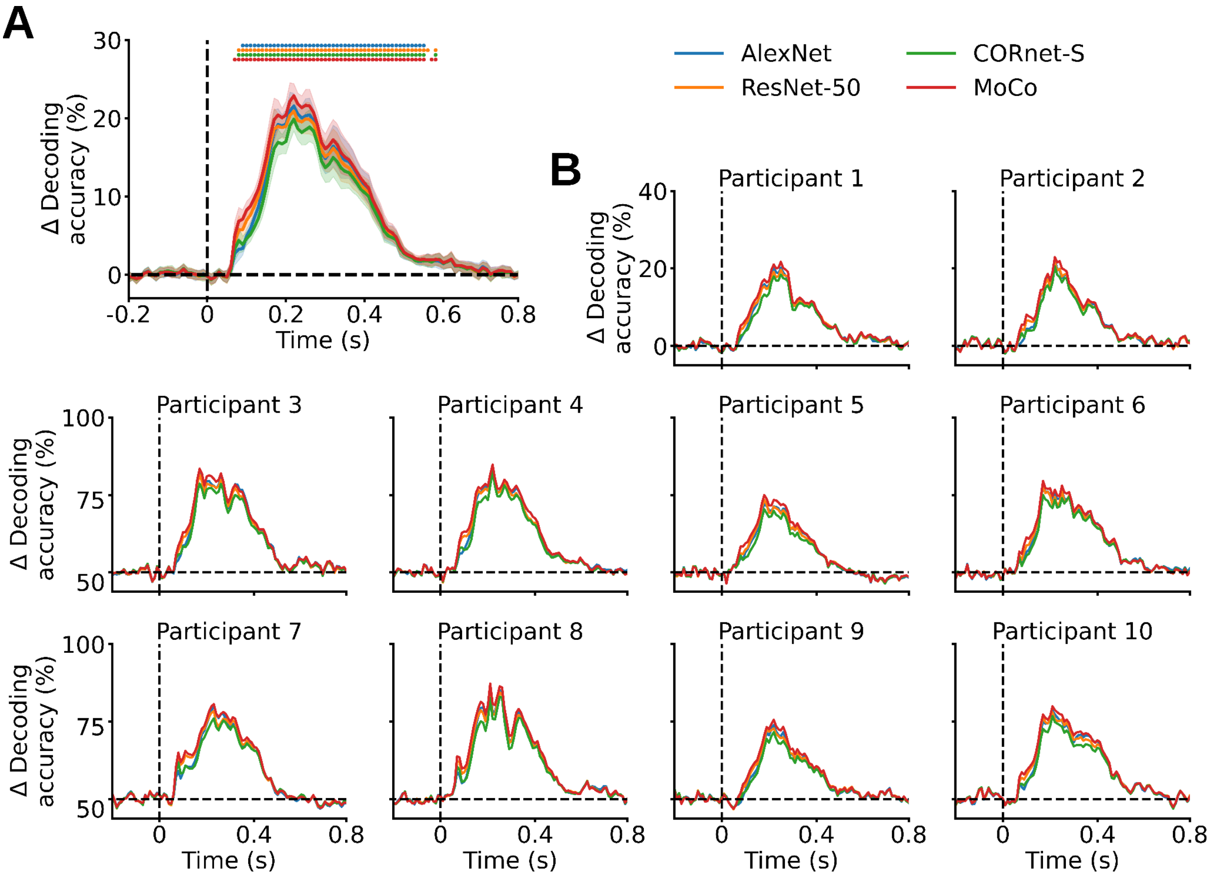


**Supplementary Figure 2.** Difference between the pairwise decoding results and the noise ceiling lower bound estimates. (**A**) The differences averaged across participants are significantly above zero between 70ms and 580ms, with peaks at 220ms ($P < 0.05$, one-sample one-sided t-test, Bonferroni-corrected). (**B**) Individual participants' results. Error margins, asterisks and black dashed lines as in **Figure 3**.

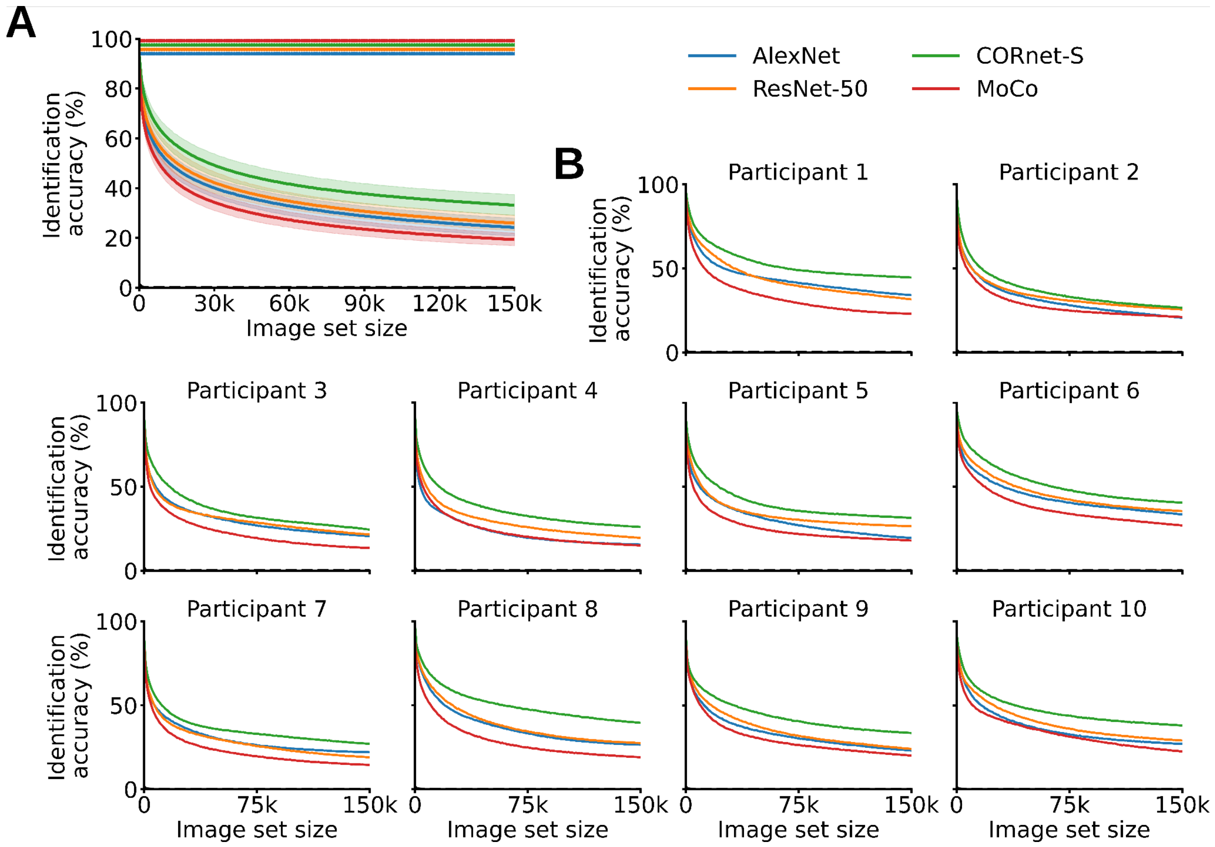


**Supplementary Figure 3.** Zero-shot identification, the three most correlated candidate image conditions. Zero-shot identification of the BioTest data using the SynTest data and the synthesized EEG visual responses to the 150,000 ILSVRC-2012 validation and test image conditions (SynImagenet), with the correct image condition falling within the three most correlated image conditions. (**A**) Zero-shot identification results averaged across participants. With a SynImagenet set size of 0 the synthesized data of AlexNet, ResNet-50, CORnet-S, MoCo significantly identify the BioTest data with accuracies of, respectively, 90.6%, 91%, 93.65%, 88.7%. ($P$ < 0.05, one-sample one-sided t-test, Bonferroni-corrected). With a SynImagenet set size of 150,000 the synthesized data of AlexNet, ResNet-50, CORnet-S, MoCo significantly identify the BioTest data with accuracies of, respectively, 24.2%, 25.95%, 33.15%, 19.35%. (**B**) Individual participants' results. Error margins and black dashed lines as in **Figure 3**. Asterisks as in **Figure 5**.

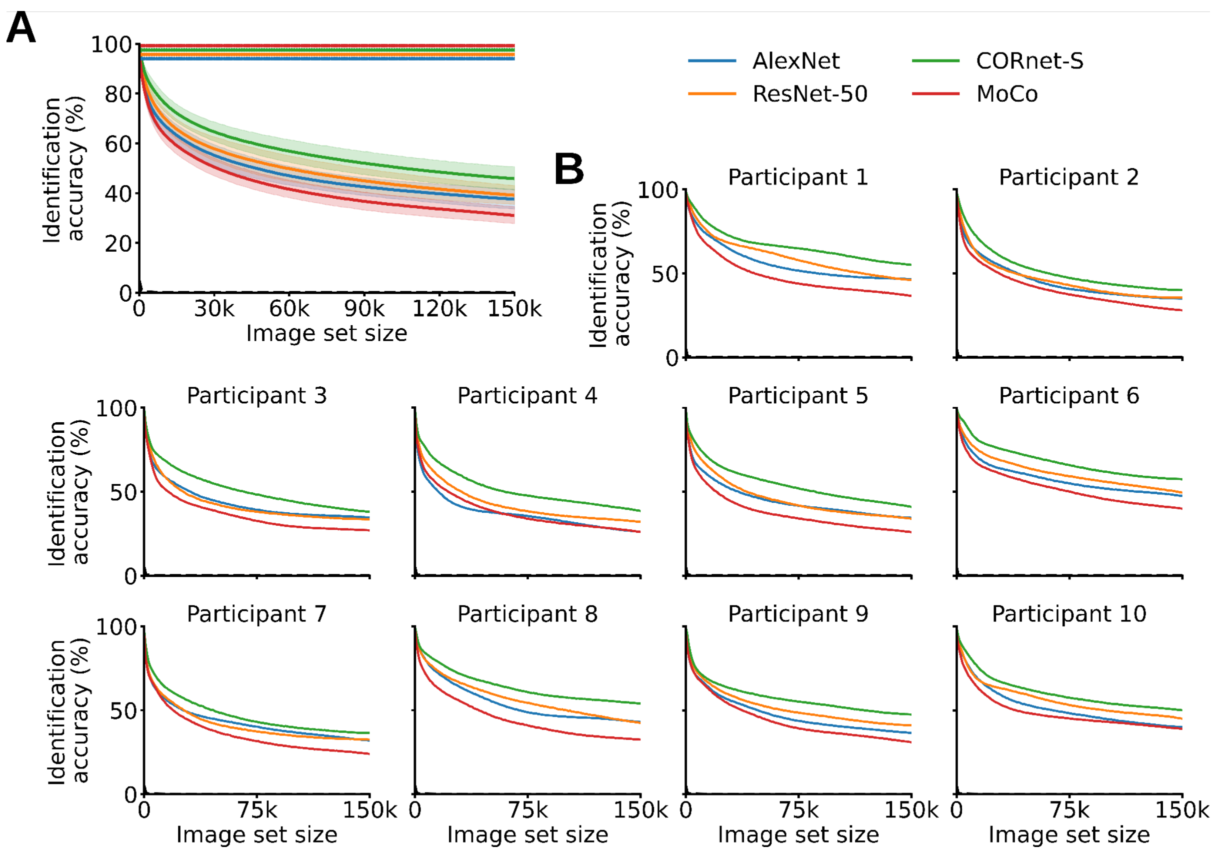


**Supplementary Figure 4.** Zero-shot identification, ten most correlated candidate image conditions. Zero-shot identification of the BioTest data using the SynTest data and the synthesized EEG visual responses to the 150,000 ILSVRC-2012 validation and test image conditions (SynImagenet), with the correct image condition falling within the ten most correlated image conditions. (**A**) Zero-shot identification results averaged across participants. With a SynImagenet set size of 0 the synthesized data of AlexNet, ResNet-50, CORnet-S, MoCo significantly identify the BioTest data with accuracies of, respectively, 97.55%, 97.7%, 99.05%, 97.05%. ($P$ < 0.05, one-sample one-sided t-test, Bonferroni-corrected). With a SynImagenet set size of 150,000 the synthesized data of AlexNet, ResNet-50, CORnet-S, MoCo significantly identify the BioTest data with accuracies of, respectively, 37.55%, 39.15%, 45.8%, 31%. (**B**) Individual participants' results. Error margins and black dashed lines as in **Figure 3**. Asterisks as in **Figure 5**.

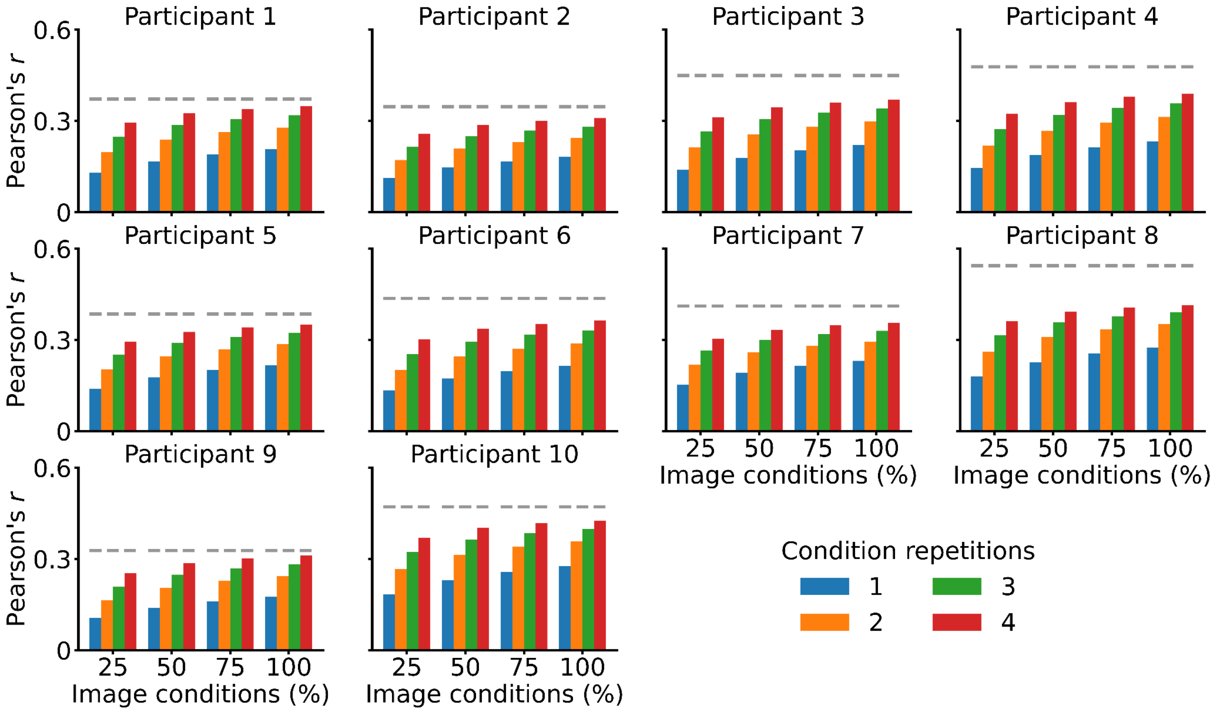


**Supplementary Figure 5.** Effect of image conditions and condition repetitions on linearizing encoding models’ prediction accuracy, individual participants’ results. Gray dashed lines as in **Figure 6**.

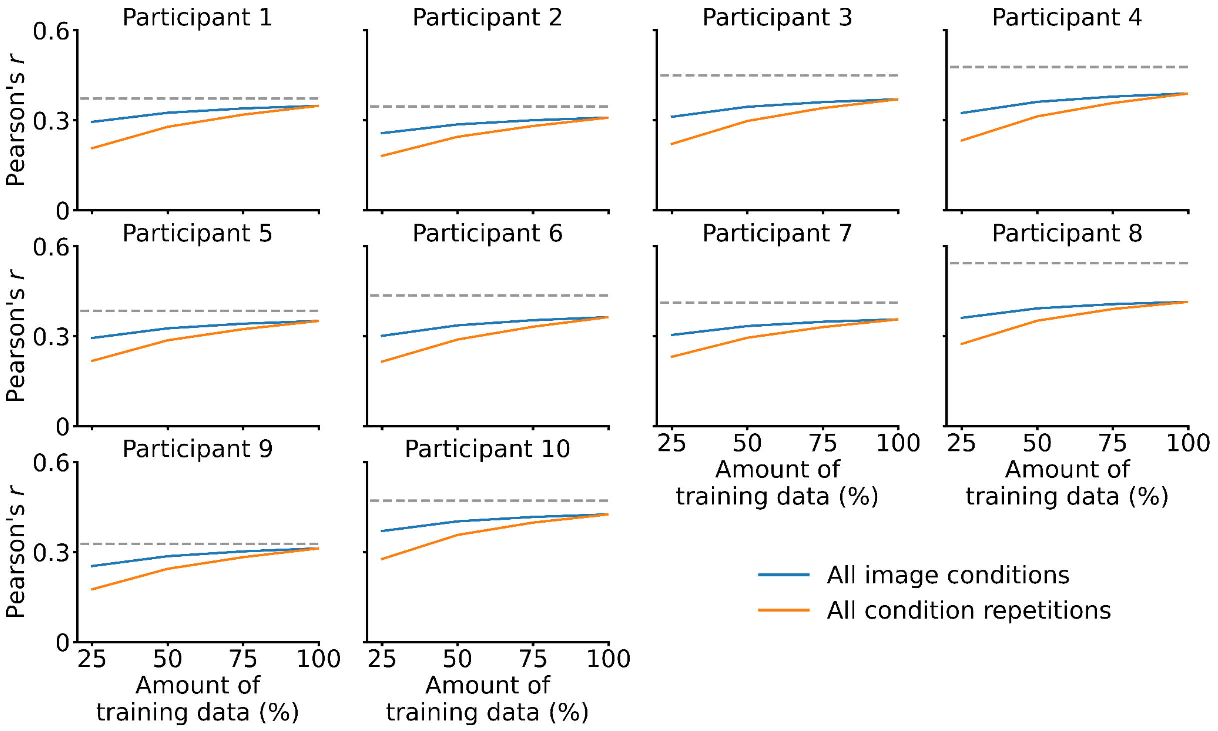


**Supplementary Figure 6.** Contribution of image conditions and condition repetitions on linearizing encoding models' prediction accuracy, individual participants' results. Gray dashed lines as in **Figure 6**.

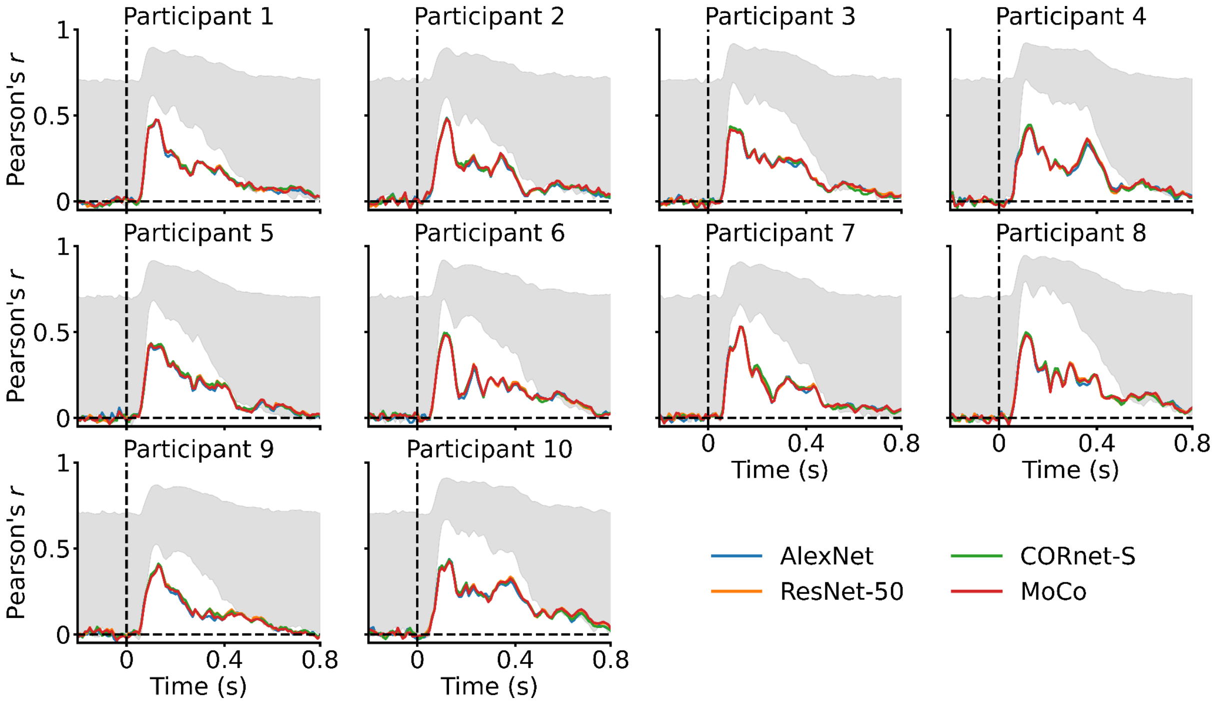


**Supplementary Figure 7.** Evaluating the prediction accuracy of linearizing encoding models which generalize to novel participants through correlation, individual participants' results. Gray areas and black dashed lines as in **Figure 3**.

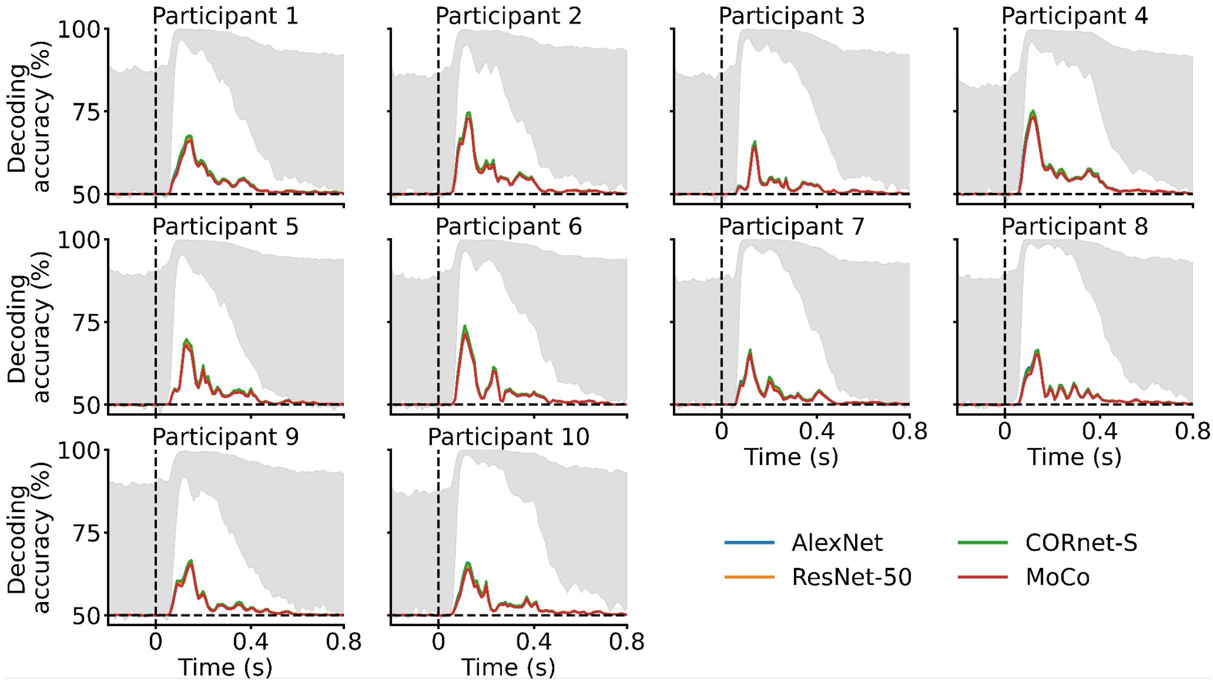


84
85 **Supplementary Figure 8.** Evaluating the prediction accuracy of linearizing encoding
86 models which generalize to novel participants through pairwise decoding, individual
87 participants' results. Gray areas and black dashed lines as in **Figure 3**.

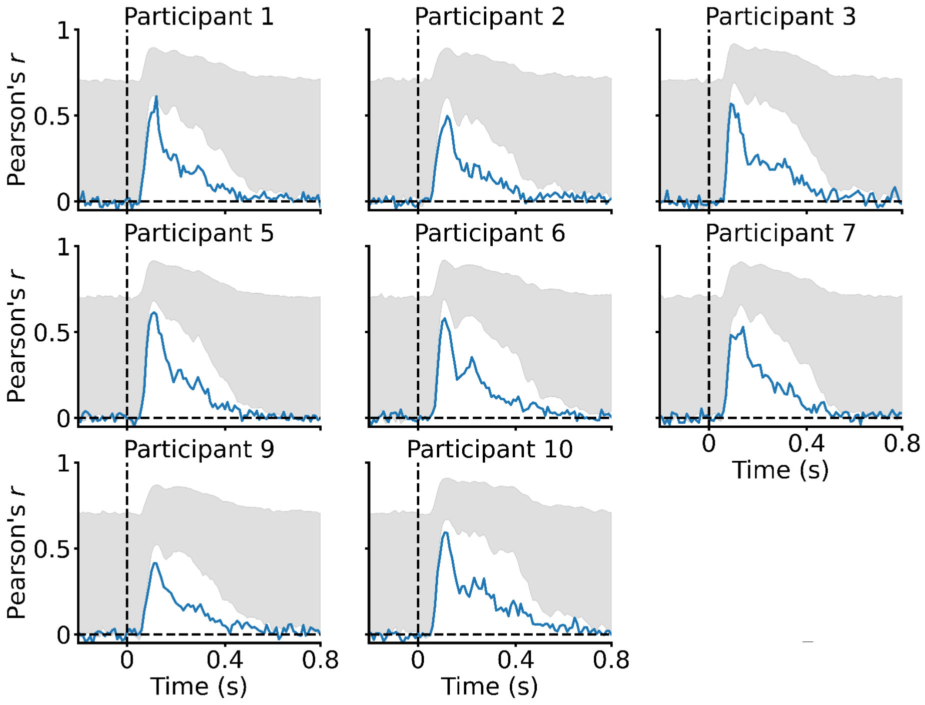


**Supplementary Figure 9.** Evaluating the end-to-end encoding models' prediction accuracy through correlation, individual participants' results. Gray areas and black dashed lines as in **Figure 3**.

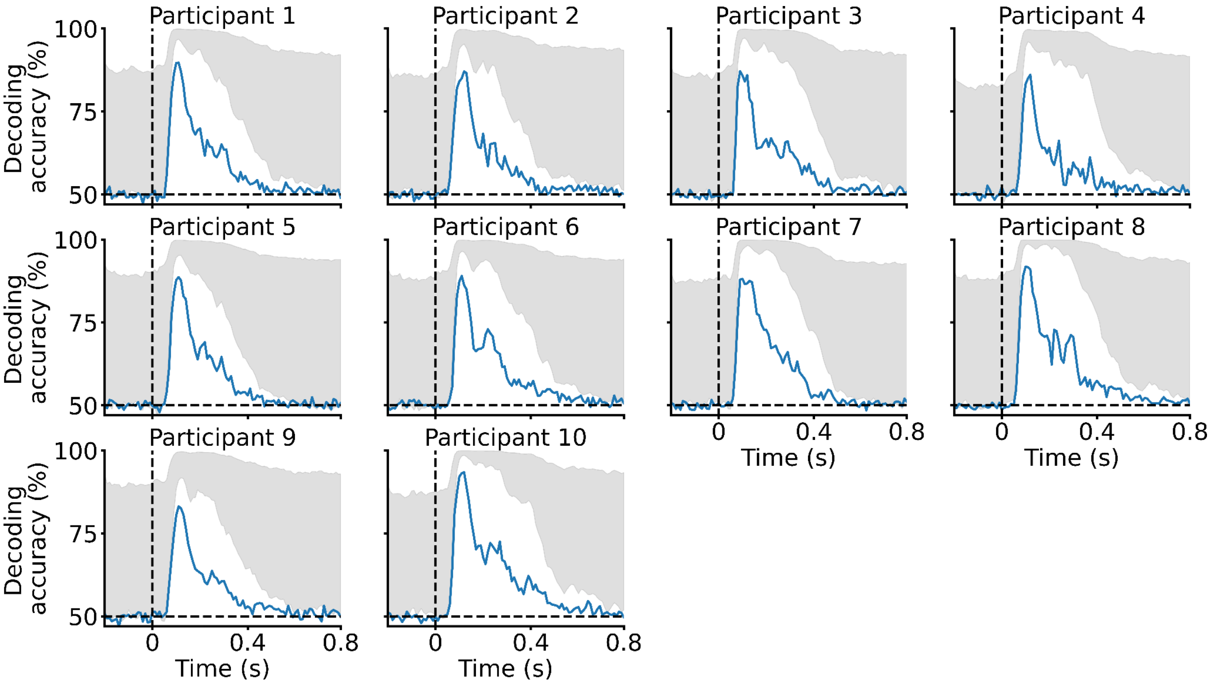


**Supplementary Figure 10.** Evaluating the end-to-end encoding models' prediction accuracy through pairwise decoding, individual participants' results. Gray areas and black dashed lines as in **Figure 3**.

| | Identification accuracy < 10% | | | | Identification accuracy < 0.5% | | | |
|---|---|---|---|---|---|---|---|---|
| | AlexNet | ResNet-50 | CORnet-S | MoCo | AlexNet | ResNet-50 | CORnet-S | MoCo |
| **Participant 1** | 2.12E+07 | 8.12E+06 | 2.65E+09 | 1.26E+06 | 3.38E+12 | 2.26E+11 | 9.46E+17 | 3.32E+09 |
| **Participant 2** | 1.03E+06 | 5.83E+06 | 3.35E+06 | 3.21E+06 | 2.20E+09 | 6.52E+11 | 4.59E+10 | 7.25E+11 |
| **Participant 3** | 1.05E+06 | 1.40E+06 | 2.61E+06 | 2.63E+05 | 3.20E+09 | 6.57E+09 | 2.75E+10 | 8.38E+07 |
| **Participant 4** | 4.35E+05 | 8.94E+05 | 2.85E+06 | 3.76E+05 | 9.49E+08 | 2.33E+09 | 3.02E+10 | 3.74E+08 |
| **Participant 5** | 6.98E+05 | 1.83E+07 | 5.18E+07 | 1.14E+06 | 5.91E+08 | 5.08E+13 | 2.33E+14 | 3.57E+10 |
| **Participant 6** | 1.84E+07 | 1.86E+07 | 3.41E+07 | 4.24E+06 | 2.41E+12 | 1.74E+12 | 4.06E+12 | 8.52E+10 |
| **Participant 7** | 2.23E+06 | 7.09E+05 | 8.31E+06 | 3.49E+05 | 7.95E+10 | 9.34E+08 | 1.21E+12 | 3.49E+08 |
| **Participant 8** | 2.11E+06 | 2.52E+06 | 4.11E+07 | 7.77E+05 | 8.70E+09 | 1.27E+10 | 7.81E+12 | 1.72E+09 |
| **Participant 9** | 1.55E+06 | 1.53E+06 | 1.18E+07 | 1.12E+06 | 5.67E+09 | 3.79E+09 | 6.23E+11 | 4.95E+09 |
| **Participant 10** | 4.27E+06 | 4.08E+06 | 1.01E+08 | 1.01E+06 | 1.16E+11 | 4.66E+10 | 2.30E+14 | 9.80E+08 |
| **Average** | 5.30E+06 | 6.20E+06 | 2.91E+08 | 1.37E+06 | 6.01E+11 | 5.35E+12 | 9.47E+16 | 8.57E+10 |

**Supplementary Table 1.** Extrapolating the zero-shot identification accuracy, three most correlated candidate image conditions. The zero-shot identification accuracy is extrapolated as a function of candidate image set sizes. The values in the table indicate the candidate image set sizes required for the identification accuracy to drop below 10% and 0.5%.

| | Identification accuracy < 10% | | | | Identification accuracy < 0.5% | | | |
|---|---|---|---|---|---|---|---|---|
| | AlexNet | ResNet-50 | CORnet-S | MoCo | AlexNet | ResNet-50 | CORnet-S | MoCo |
| Participant 1 | 7.76E+08 | 2.11E+07 | 9.75E+08 | 3.27E+07 | 1.60E+16 | 3.34E+11 | 4.16E+15 | 7.03E+12 |
| Participant 2 | 1.81E+07 | 9.79E+06 | 2.99E+07 | 2.38E+06 | 2.04E+12 | 2.33E+11 | 3.09E+12 | 6.30E+09 |
| Participant 3 | 4.99E+07 | 5.18E+07 | 9.74E+06 | 2.93E+06 | 7.16E+13 | 1.15E+14 | 1.06E+11 | 3.08E+10 |
| Participant 4 | 2.65E+06 | 1.42E+07 | 3.32E+07 | 2.55E+06 | 1.42E+10 | 1.65E+12 | 4.05E+12 | 1.44E+10 |
| Participant 5 | 1.96E+07 | 9.22E+06 | 1.77E+07 | 2.54E+06 | 2.34E+12 | 2.16E+11 | 3.79E+11 | 1.47E+10 |
| Participant 6 | 3.63E+08 | 2.53E+08 | 7.38E+08 | 1.45E+07 | 1.09E+15 | 2.28E+14 | 1.67E+15 | 2.70E+11 |
| Participant 7 | 8.34E+06 | 2.51E+07 | 1.70E+07 | 1.53E+06 | 2.24E+11 | 1.36E+13 | 1.13E+12 | 4.02E+09 |
| Participant 8 | 1.41E+08 | 1.73E+07 | 1.55E+09 | 4.30E+06 | 1.90E+14 | 2.73E+11 | 2.16E+16 | 2.40E+10 |
| Participant 9 | 1.99E+07 | 5.35E+07 | 3.48E+08 | 5.37E+06 | 1.60E+12 | 1.40E+13 | 9.77E+14 | 5.83E+10 |
| Participant 10 | 3.43E+07 | 1.18E+08 | 6.37E+08 | 3.11E+08 | 4.12E+12 | 6.30E+13 | 3.45E+15 | 4.96E+15 |
| Average | 1.43E+08 | 5.73E+07 | 4.35E+08 | 3.80E+07 | 1.73E+15 | 4.37E+13 | 3.19E+15 | 4.97E+14 |

**Supplementary Table 2.** Extrapolating the zero-shot identification accuracy, ten most correlated candidate image conditions. The zero-shot identification accuracy is extrapolated as a function of candidate image set sizes. The values in the table indicate the candidate image set sizes required for the identification accuracy to drop below 10% and 0.5%.
